# Age-dependent tumor-immune interactions underlie immunotherapy response in pediatric cancer

**DOI:** 10.1101/2025.07.16.663652

**Authors:** Omar Elaskalani, Zahra Abbas, Sébastien Malinge, Merridee A. Wouters, Jenny Truong, Iley M. Johnson, Jorren Kuster, Alexander Nassar, Allison Wan, Hannah Smolders, Hilary Hii, Alison M. McDonnell, Isaac Popal, Grace A. Chua, Vivien Nguyen, Joyce Oommen, Sajla Singh, Febriana Ajelie, Kunjal Panchal, Annabel Short, Meegan Howlett, Rachael M. Zemek, Ben Wylie, Jonathan Chee, Bree Foley, Claudia L. Kleinman, Nada Jabado, Timothy N. Phoenix, Misty R. Jenkins, Nicholas G. Gottardo, Terrance G. Johns, Rishi S. Kotecha, Laurence C. Cheung, Timo Lassmann, Raelene Endersby, W. Joost Lesterhuis

## Abstract

Pediatric cancers originate in rapidly growing tissues within the context of a developing host. However, the interactions between cancer cells and the developing immune system are incompletely understood. Here, we established a suite of pediatric syngeneic mouse cancer models across diverse anatomical sites and compared their tumor immune microenvironment with that in adult mice. Tumors in pediatric mice exhibited significantly accelerated growth and diminished leukocyte infiltration, dominated by naïve-like PD-1^low^/CD8^+^ T cells, and proliferative MHCII^low^/PD-L1^hi^/CD86^low^ macrophages. Tumor-infiltrating leukocytes in pediatric mice were enriched for MYC targets, which was also observed in pediatric patient samples. Furthermore, pediatric mice displayed poor responses to anti-PD-1/PD-L1 or bispecific T cell engager antibodies, which could be reversed by inducing a proinflammatory microenvironment via MYC inhibition or inducing macrophage polarization to an MHCII^hi^ phenotype. These findings underscore the significant influence of young age on cancer immune responses and reveal potential new therapeutic opportunities for pediatric cancers.

**HIGHLIGHTS:**

- Allograft tumors exhibit markedly accelerated growth in pediatric hosts compared to adults.
- Tumors growing in pediatric mice have reduced leukocyte infiltration, dominated by naïve-like PD-1^low^/CD8^+^ T cells, and MHCII^low^/M2-like macrophages.
- Enrichment of MYC target genes is observed in pediatric mouse tumors and confirmed in primary patient tumor samples.
- Pediatric mice display reduced response to anti-PD-1/PD-L1 and BiTE immunotherapy, which can be reversed by remodeling the TIME, using either MYC inhibition or macrophage polarization.

## INTRODUCTION

The tumor immune microenvironment (TIME) significantly influences tumor growth and metastasis, making it an important target for therapy^1^. However, in contrast to the extensive data available from adult patients, our understanding of the TIME in pediatric cancers is limited, impeding the development of effective immunotherapeutic strategies for childhood cancers^2,3^. Significant progress has been made with T cell-directed immunotherapies in children with leukemia, such as chimeric antigen receptor (CAR)-T cells and bispecific T cell engager (BiTE) antibodies^4,5^. In contrast, therapeutic targeting of the TIME has shown minimal success in other pediatric cancers^6^. Whilst immune checkpoint therapy (ICT) targeting programmed cell death protein 1 (PD-1) or its ligand PD-L1 have shown substantial efficacy in adult cancers, these therapies have yielded disappointing outcomes in nearly all pediatric solid cancers^7,8^.

Recent single-cell RNA sequencing studies have identified crucial differences in the anti-cancer immune response between children and adults, indicating that the pediatric TIME is more immunosuppressive compared to the adult TIME^9–12^. However, pediatric cancers are often different from adult cancers in terms of their cell of origin, mutational burden, and in their relative frequency of oncogenic drivers^13,14^, making it challenging to determine whether any observed differences in the TIME and the response to immunotherapy between pediatric and adult patients are host—derived or cancer cell—intrinsic. This gap in understanding the determinants of the pediatric TIME impedes the development of effective immunotherapy for childhood cancers^6^.

Preclinical cancer studies invariably involve adult mice, even in pediatric cancer research^15–20^. This practice may overlook crucial developmental influences on pediatric cancer cell biology, particularly when identifying and testing treatments that target the TIME rather than cancer cell-intrinsic vulnerabilities. Here, we set out to identify age—specific, host-derived mechanisms underlying the anti-cancer immune responses by developing a suite of syngeneic allograft cancer mouse models of pediatric age. We comprehensively characterized the TIME and response to immunotherapy in these pediatric models, in comparison with the same cancer models in adult mice. Our findings reveal key age-related differences, which we confirmed to be consistent with human pediatric tumor data across different age categories. Moreover, therapeutic modulation of the TIME improved cancer control in young mice, revealing valuable new treatment opportunities.

## RESULTS

### Tumor growth is accelerated in pediatric hosts

To explore the impact of age-related host immunobiology on pediatric cancers, we selected murine cancer models that reproducibly form tumors upon implantation into immune-competent strains and compared tumor growth kinetics in pediatric or adult mice. This approach allowed us to specifically examine the influence of age whilst keeping other variables consistent, such as host genetics, tumor type, cancer cell genetics, and environmental influences, in an immunocompetent *in vivo* setting. We employed five distinct models representing three common pediatric cancer types: leukemia, brain cancer, and sarcoma (**Fig. 1A**). These included group 3 medulloblastoma (G3MB) cells driven by *Myc* overexpression and dominant negative p53 (Myc/p53^DD^) injected orthotopically in the cerebellum (**Fig. 1B-C**)^21^; a rhabdomyosarcoma (RMS) cell line derived from *Mt1:Hgf*^Tg^;*Tp53*^+/-^ mice^22^ (M3-9-M), either injected subcutaneously (**Fig. 1D**); or orthotopically in the gastrocnemius muscle (**Fig. 1E-F**); the carcinogen-induced fibrosarcomas MCA205^23^ and WEHI164^19^, injected subcutaneously (**Fig. 1D**); and murine B-cell acute lymphoblastic leukemia (B-ALL) driven by *BCR::ABL* fusion (PER-M60)^20^ injected intravenously (**Fig. 1G**). These models encompass both highly mutated cancers resembling adult malignancies (MCA205 and WEHI164 fibrosarcomas), and cancers initiated by a few genetic drivers, closely mimicking pediatric disease (Myc/p53^DD^ G3MB, M3-9-M RMS, PER-M60 B-ALL, **Fig. S1A**). To compare ages, pediatric mice were implanted at postnatal day (P) 1-4 (pups), while mice implanted at 12-14 weeks of age were considered adult. Across all models, we observed markedly accelerated cancer growth in pediatric mice compared to adult mice when the same number of cells were implanted (**Fig. 1B-H**). The data were further confirmed in additional models which included: H3.1^K27M^ diffuse midline glioma (DMG) cells implanted in the brain stem^24^, MLL-AF9 acute myeloid leukemia (PER-M5) cells implanted intravenously, and a neuroblastoma cell line derived from *TH:MYCN* transgenic mice (9464D)^25^ implanted subcutaneously (**Fig. S1B-F**). To test whether a difference in body or organ size could explain the observed difference in tumor growth, we implanted ten-fold more M3-9-M RMS cells into adult mice subcutaneously, or 80-fold more PER-M60 B-ALL cells intravenously. Still, the tumor growth in pediatric mice was significantly faster (**Fig. S1G-H**). These effects were consistent across different mouse strains (C57BL/6J and BALB/c), anatomical locations (brain, muscle, bone marrow or subcutaneous tissue), and sexes (**Fig. S1I-J**).

**Figure 1.**
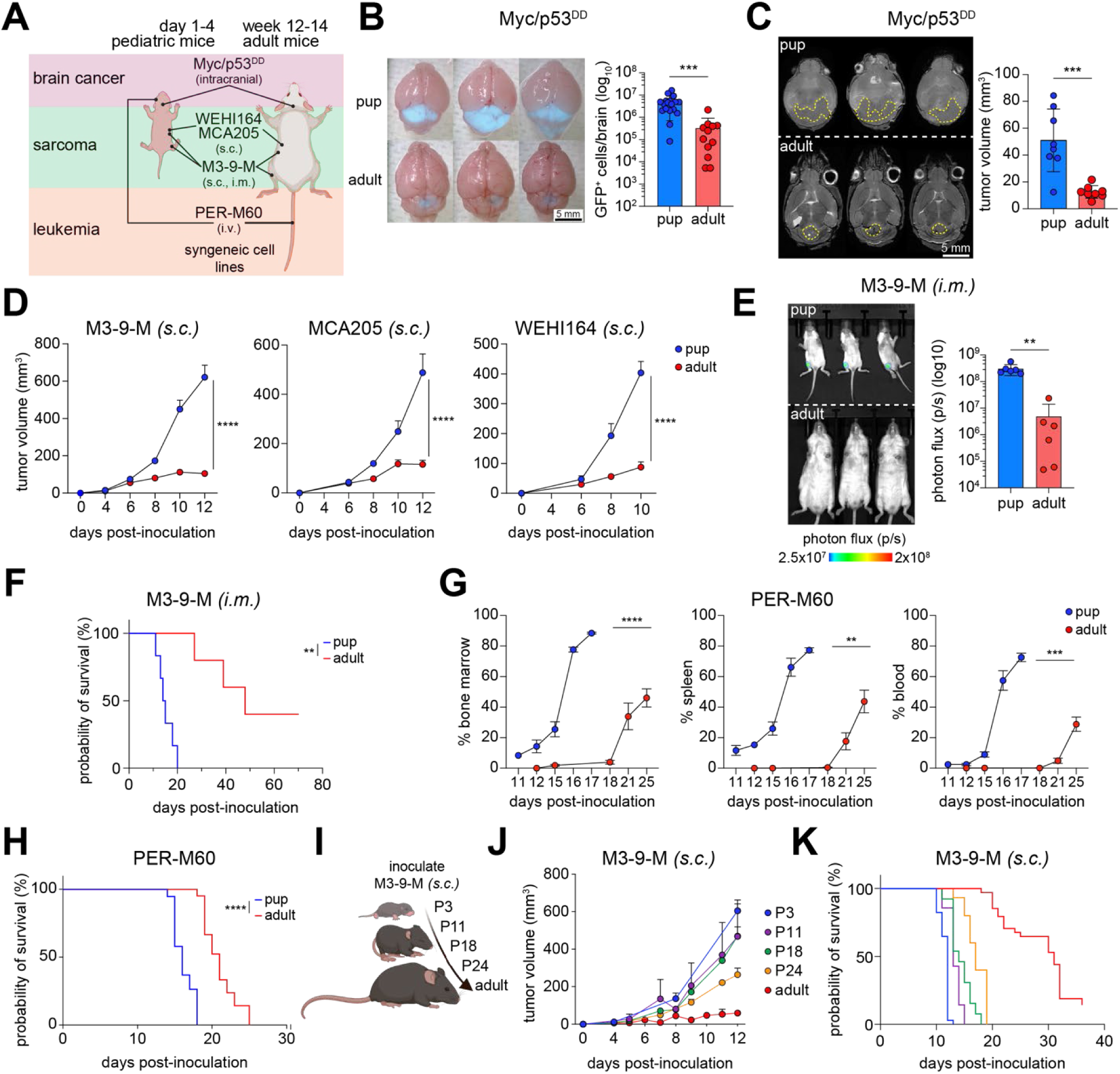
Tumor growth is accelerated in pediatric hosts. (A) Experimental setup depicting implantation of tumor cells into mice via intracranial, subcutaneous (*s.c.),* intramuscular (*i.m*.) or intravenous (*i.v.*) routes. (B) Representative images of pup and adult brains 10 days post—inoculation of GFP^+^ Myc/p53^DD^ G3MB cells *(left)* and enumeration of GFP^+^ cells in the whole brain (n = 18 pup, n = 12 adult) *(right)*. (C) Representative T2-weighted MRI images of pup and adult brains 10 days-post—inoculation of Myc/p53^DD^ G3MB cells *(left, tumor margins in yellow)*, and tumor volumes (n = 8 per age) *(right)*. (D) Tumor volumes of sarcoma cell lines implanted subcutaneously in the right flank of pups and adults (M3-9-M: n = 30 pup, n = 27 adult; MCA205: n = 20 pup, n = 22 adult; WEHI164 n = 24 pup, n = 19 adult). (E) Representative images *(left)* and quantified bioluminescent flux *(right)* of M3-9-M RMS implanted intramuscularly *(i.m.)* into the gastrocnemius of pups (n = 7) or adult mice (n = 12) 7 days post-inoculation. (F) Survival of pups (n = 6) and adults (n = 6) bearing orthotopic M3—9—M RMS. (G) Percentage of intravenously inoculated PER-M60 B-ALL cells in bone marrow, spleen, and blood of pups and adult mice (n ≥ 5/group/timepoint). (H) Survival of pups (n = 19) and adult mice (n = 21) bearing PER-M60 B-ALL. (I) Experimental schema for J-K. (J) Tumor volume measurements of M3-9-M RMS over time following inoculation into P3 (n=8), P11 (n = 8), P18 (n = 11), P24 (n = 13), and adult mice (n = 13). (K) Survival of pediatric P3 (n = 34), P11 (n = 8), P18 (n = 13), P24 (n = 15), and adult mice (n = 34) bearing M3-9-M RMS. Linear mixed-effect modelling revealed tumor growth rate decreased with age (est. fixed effect of time:age interaction = -0.0028; t = -9.88). Data presented as mean ± SEM. Significance was calculated using an unpaired Student’s t-test (B, C, E), mixed-effects models from three independent experiments (D, G, J) or Log-rank (Mantel-Cox) test from two independent experiments (F, H). * p < 0.05, ** p < 0.01, *** p < 0.001, **** p < 0.0001.

To further investigate the effect of age on tumor growth, mice were implanted with either M3-9-M RMS, Myc/p53^DD^ G3MB, or PER-M60 B-ALL cells at increasing ages, which revealed a graded, age-dependent effect on tumor growth (**Fig. 1I-K, Fig. S1K-M**). Linear mixed-effect modelling revealed tumor growth rate decreased with age (**Fig. 1I-K**). These results demonstrate that a young host provides a permissive environment for enhanced tumor growth.

### Age shapes tumor leukocyte infiltration

Immune cell profiles in peripheral blood and tissues of children are distinct from adults, both in healthy conditions and during infectious diseases^26–29^. We therefore sought to understand how pediatric age influences the nature of tumor-infiltrating leukocytes (TILs) in mice. Multispectral flow cytometric analysis of tumor suspensions from pediatric and adult mice inoculated with identical cancer cells (Myc/p53^DD^ G3MB, M3-9-M RMS or PER-M60 B-ALL) revealed that tumors in pediatric mice had significantly fewer CD45^+^ TILs (**Fig. 2A-C, Fig. S2A**). TILs in adult mice exhibited a higher proportion of CD4^+^ and CD8^+^ T cells, natural killer (NK) cells (NK1.1^+^/CD3^-^), and dendritic cells (DCs, CD11b^+^/MHCII^hi^/CD11C^hi^). In contrast, the TIL populations in pediatric mice were predominantly comprised of tumor-associated macrophages (TAMs, F4/80^hi^/Ly6C^-^) across all models, and monocytes (Ly6C^hi^/F4/80^int^) in all models except PER-M60 B-ALL (**Fig. 2A-C**, gating strategy is available in additional information). Except for CD4⁺ T cells and monocytes, these differences in immune cell frequency were also observed between healthy pediatric and adult control mice (**Fig. S2B**). Since increased tumor mutational burden is associated with increased TILs in some adult cancers^30^. We sought to examine whether implanting highly mutated tumors in pediatric mice would similarly increase TILs or also result in an age-dependent reduction. Using the carcinogen-induced, highly mutated MCA205 fibrosarcoma implanted into both pediatric and adult mice, we observed that tumors in pediatric mice also contained fewer TILs compared to adult mice (**Fig. S2C**). While there were no differences in TAM abundance, MCA205 fibrosarcoma in pediatric mice had fewer CD8^+^ T cells, NK cells and DCs and more monocytes compared to adult mice (**Fig. S2C**). Altogether, this data suggest that age-related differences in TILs are observed despite intrinsic genetic differences of the cancer cells tested.

**Figure 2.**
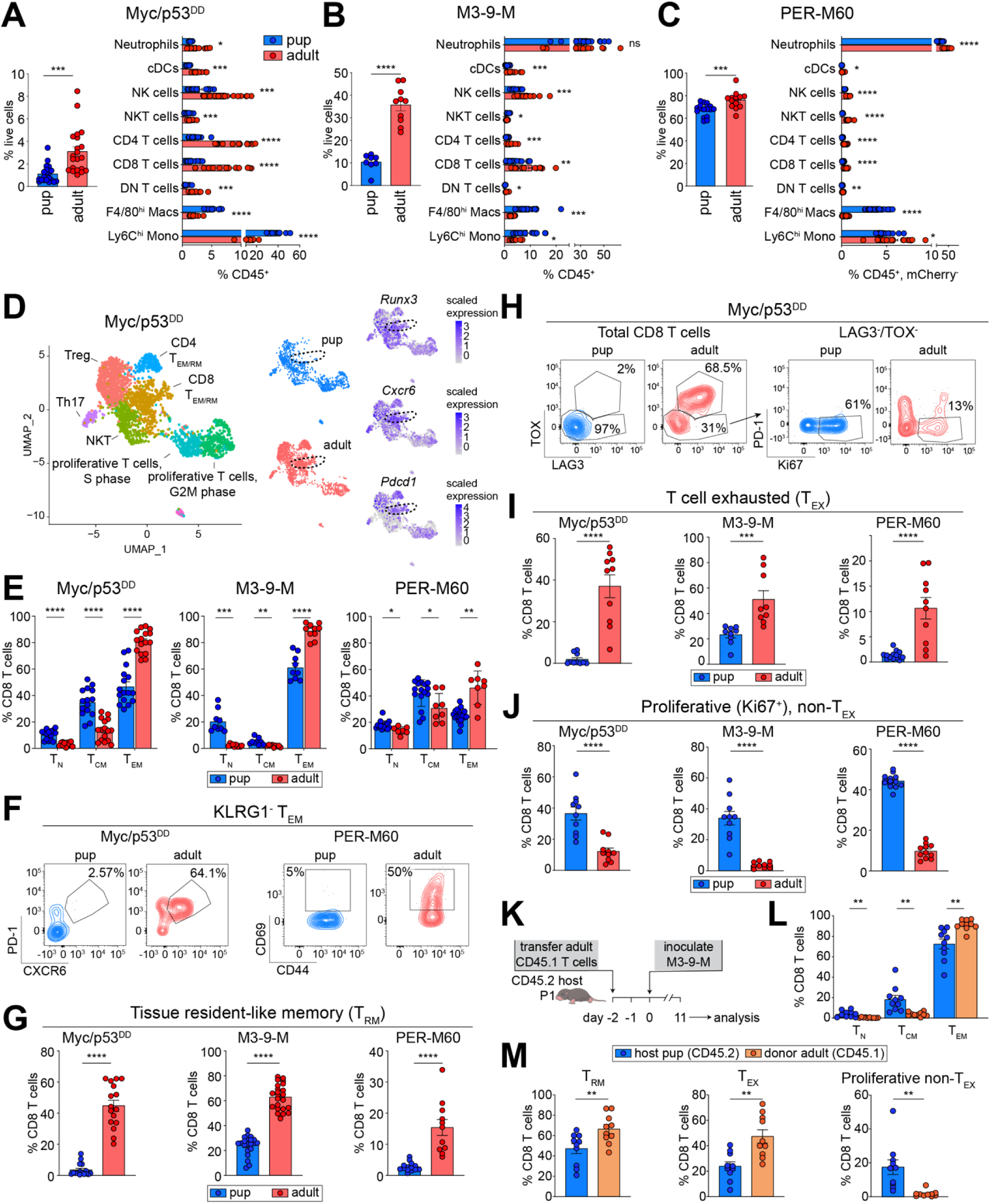
Tumor infiltration of leukocytes and CD8+ T cell fate is age dependent. (A-) Frequency of CD45^+^ tumor-infiltrating leukocytes (TILs) as a percentage of live cells *(left)* and abundance of different immune cells as a percentage of total CD45^+^ cells (*right*) in (A) Myc/p53^DD^ G3MB and (B) M3-9-M RMS, or (C) as a percentage of CD45^+^ cells in the bone marrow after excluding PER-M60 B-ALL (mCherry^+^) cells *(right)*. (D) UMAP of scRNAseq data showing T cells in Myc/p53^DD^ G3MB in pups and adult mice *(left, middle)*, and the expression pattern of *Runx3*, *Pdcd1* and *Cxcr6 (right)*. (E) Frequency of T_N_, T_CM_ and T_EM_ in CD8^+^ T cells in pups and adult mice 11-15 days after cancer cell inoculation. (F) Representative flow cytometry plots of CD8^+^ T_RM_ in solid tumor models (PD-1^+^CXCR6^+^KLRG1^-^T_EM_) *(left, Myc/p53^DD^ shown)* and in PER-M60 B-ALL (CD69^+^KLRG1^-^T_EM_) *(right)* and (G) the frequency of T_RM_ in pups and adult mice. (H) Representative flow cytometry plots from Myc/p53^DD^ tumors of CD8^+^ T_EX_ (LAG3^+^TOX^+^) *(left)* and proliferative non-T_EX_ (Ki67^+^PD-1^-^LAG3^-^TOX^-^) *(right)* and (I) the frequency of T_EX_ and (J) proliferative non-T_EX_ in CD8^+^ T cells. (K) Adoptive T cell transfer schema. (L) Frequency of T_N_, T_CM_, and T_EM_ and (M) frequency of T_RM_, T_EX_, proliferative non-T_EX_ CD8^+^ T cells of intratumoral host pup and donor adult cells as a percentage of total host and donor CD8^+^ T cells, respectively. Data presented as mean ± SEM. Sample sizes: n ≥ 8 per group (A-C and E-M), from ≥ 2 independent experiments, n = 4 per group (D). Groups were compared using unpaired Student’s t-test with Holm-Šídák correction for multiple comparisons (A-C, E, L) or unpaired Student’s t-test (G, I, J, M). * p < 0.05, ** p < 0.01, *** p < 0.001, **** p < 0.0001.

### Age affects intratumoral CD8^+^ T cell differentiation, resident memory formation, exhaustion and proliferation

We next sought to determine if there are specific age-dependent differences in tumor-infiltrating immune cell phenotypes. We focused on CD8^+^ T cells and TAMs due to their consistent age-dependent differences within TILs across all models, in addition to their known role in immunotherapy response in clinical trials across pediatric and adult cancers^31^. We performed single cell RNA sequencing (scRNAseq) on Myc/p53^DD^ G3MB and M3-9-M tumors and bone marrow from the PER-M60 B-ALL model. After standard clustering^32^, T cells were analyzed separately and subclustered. The higher number of recovered T cells in Myc/p53^DD^ G3MB (owing to enrichment for CD45^+^ cells during sample preparation) allowed for the identification of 7 subpopulations (**Fig. 2D, Table S1**). We noted that pediatric tumors had relatively fewer CD8^+^ T cells that express markers associated with long-term resident memory (*Cxcr6, Runx3, Itga1* and *CD69*)^33^ and exhaustion (*Pdcd1, Lag3* and *Tox*)^34^ (**Fig. 2D, Fig. S3A, Table S1**), which were consistent in the M3-9-M RMS model (**Fig. S3B-C, Table S1**). To confirm and expand on these differences, we analyzed intratumoral CD8^+^ T cells using spectral flow cytometry. We started by investigating the impact of age on markers of differentiation, resident memory, exhaustion, and proliferation. Across Myc/p53^DD^ G3MB, M3-9-M RMS, and PER-M60 B-ALL models, we found that intratumoral pediatric CD8^+^ T cells displayed a less differentiated, naïve-like (T_N_, CD62L^+^/CD44^-^) and central memory (T_CM_, CD62L^+^/CD44^+^) phenotype, at the expense of effector memory (T_EM_, CD62L^-^/CD44^+^) differentiation (**Fig. 2E**). Within the CD8^+^ T_EM_ population, pediatric tumors contained significantly fewer resident-like memory T cells (T_RM_) expressing CXCR6, RUNX3, CD69, CD49a and the immune checkpoint PD-1^35–38^ compared to adults (**Fig. 2F-G, Fig. S3D-F**). To distinguish tumor-resident CD8^+^ T cells from circulating ones, we injected mice intravenously with a PE-conjugated CD45 antibody. This allowed us to differentiate blood-borne CD8+ T cells (PE^+^) from tumor-resident CD8+ T cells (PE^−^)^39,40^. In both the M3-9-M RMS and PER-M60 B-ALL models, adult mice exhibited a higher proportion of CD8 T_RM_ (characterized as PE^−^/CD62L^−^/KLRG1^−^/CD44^+^ and CXCR6^+^ or CD69^+^) compared to pups (**Fig. S3G-J**). Furthermore, pediatric CD8^+^ T cells were generally less exhausted (T_EX_, LAG3^+^/TOX^+^/PD-1^+^)^41^ and instead were enriched for subsets that could be further refined as non-exhausted proliferative (PD-1^-^/Ki67^+^)^42^ (**Fig. 2H-J**). This unique phenotype of intratumoral pediatric CD8^+^ T cells, relative to adults, was not influenced by tumor burden (**Fig. S3K-M**), and persisted in highly mutated sarcoma (**Fig. S3N-O**). Although pediatric and adult CD8⁺ T cells differed in their effector memory subset under healthy conditions (**Fig. S4A**), no baseline discrepancies were observed in exhaustion subset (**Fig. S4A**). This is consistent with the known upregulation of exhaustion markers under prolonged antigenic stimulation^33^.

We next questioned whether the pediatric microenvironment drives these age—associated changes observed within tumor-infiltrating T cells. To test this, we adoptively transferred magnetically enriched splenic T cells (CD3^+^) from healthy, non-tumor bearing CD45.1^+^ adult mice into CD45.2^+^ pediatric mice prior to tumor inoculation and analyzed tumors 10-13 days post cancer cell implant. The transfer of adult T cells did not slow tumor progression in pediatric mice (**Fig. S4B**). Importantly, donor adult CD8^+^ T cells did not adopt a pediatric-like phenotype. Instead, they retained their capacity for differentiation, showing higher proportions of T_RM_ and T_EX_, with non-T_EX_ cells being less proliferative compared to endogenous pediatric CD8^+^ T cells in M3-9-M RMS (**Fig. 2K-M**), Myc/p53^DD^ G3MB and PER-M60 B-ALL (**Fig. S4C-J**). To further confirm this observation, we transferred flow-sorted naïve (CD62L^hi^/CD44^low^) adult CD45.1⁺ T cells into pediatric mice prior to tumor implantation. Similar to total adult T cells, naïve adult-derived T cells preserved their adult differentiation characteristics, preferentially forming memory and exhaustion subsets, and exhibiting reduced proliferation compared to pediatric-derived endogenous CD8⁺ T cells (**Fig. S4K-N**). This suggests that the developmental origin (pediatric or adult) of CD8^+^ T cells drives their intratumoral phenotype.

### Early life tumor-associated macrophages exhibit an MHCII^low^/M2-like phenotype

TAMs were more abundant in tumors from pediatric mice compared to adults (**Fig. 2A-C**). As TAMs can mediate T cell responses^43^ and are abundant in pediatric patient tumor samples^44–46^, we assessed their age-associated phenotype using scRNAseq. TAMs and monocytes were subclustered revealing 8 subpopulations in Myc/p53^DD^ G3MB, 8 subpopulations in M3-9-M RMS, and 10 subpopulations in PER-M60 B-ALL (**Fig. S5A**). These data revealed age- and context-dependent differences in abundance and differentiation state of myeloid cells across the three different cancer types (**Table S1**). Our combined analysis indicated that the most consistent age-dependent difference across multiple TAM (and monocyte) populations was the reduced expression of MHCII genes in pups (**Fig. 3A**). Results were validated in scRNAseq data from patients with pediatric B-ALL^47^, AML^48^ and Sonic Hedgehog medulloblastoma (SHH MB)^49^, showing lower expression of MHCII genes in all myeloid cells in younger children (less than 4y in B-ALL and AML, and 2y in SHH MB), compared to older patients (above 14y) (**Fig. 3B, Fig. S5B**). These results were also orthogonally validated across five different models using flow cytometry, demonstrating that MHCII^+^ TAMs and monocytes are less abundant in the pediatric setting (**Fig. 3C-D**), both under healthy conditions and during cancer progression (**Fig. S5C**). Notably, these pediatric MHCII^low^ TAMs were more proliferative (Ki67^+^), expressed the immune checkpoint PD-L1, and had low-level expression of the co-stimulatory molecule CD86 (**Fig. 3E-F**) compared to adults. Additionally, albeit model-dependent, the MHCII^low^ TAMs in pediatric mouse tumors also expressed M2-like markers associated with immune suppression^50–52^, including arginase 1 (ARG1), the mannose receptor CD206, folate receptor 2 (FOLR2), CD93, complement receptor of immunoglobulin superfamily (CRIg) and T cell membrane protein 4 (TIM-4) (in M3-9-M RMS and PER-M60 B-ALL) (**Fig. S5D-E**). To rule out the possibility that the higher frequency of MHCII^low^/M2-like TAMs in pediatric mouse tumors was driven by their larger size^53^ (∼600 mm^3^, compared with ∼200mm^3^ in adult mice), we compared TAMs in pediatric hosts with size-matched tumors from adult mice (**Fig. 3G**). Regardless of the tumor size, pediatric TAMs showed lower MHCII and were more M2-like (PD-L1^+^/CD206^+^/ARG1^+^) compared to large adult tumors, indicating that age is a major determinant of TAM phenotype in pediatric mice (**Fig. 3H-I**). Collectively, these data suggest that TAMs are skewed towards an M2-like immunosuppressive phenotype in early life.

**Figure 3:**
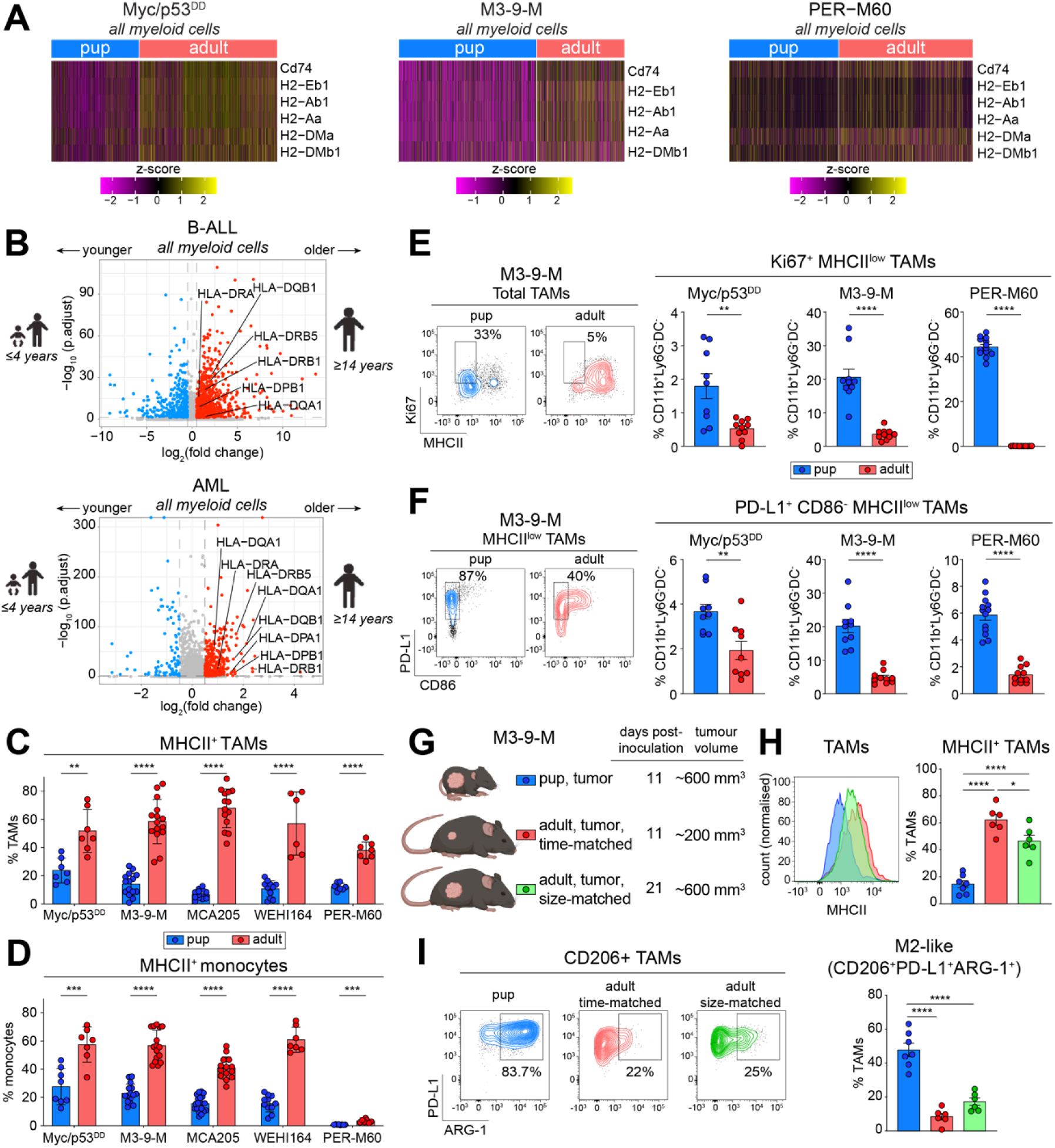
Early-life tumor-associated macrophages are MHCII^low^/M2-like. (A) Scaled expression of MHCII genes in myeloid cells in the indicated murine model. (B) Volcano plots showing differential expression of MHCII genes in myeloid cells from bone marrow of B-ALL (*top*) and AML (*bottom*) patients. (C-D) Frequency of MHCII^+^ (C) TAMs and (D) monocytes as a proportion of total TAMs and monocytes, respectively, in the indicated tumor models. (E-F) Representative flow cytometry plots *(left, from M3-9-M)* and frequency *(right)* of (E) Ki67^+^MHCII^low^ TAMs as a proportion of the parent population (CD11b^+^Ly6G^-^DCs^-^) or (F) PD-L1^+^CD86^+^MHCII^low^ TAMs as a proportion of CD11b^+^Ly6G^-^ DCs^-^ in pups and adult mice of the indicated tumor models. (G) Schema for H-I. (H) Representative histogram (*left*) and frequency (*right*) of MHCII positivity in TAMs from pups and adult mice bearing M3-9-M RMS. (I) Representative flow cytometry plots (*left*) and frequency (*right*) of M2-like (CD206^+^PD-L1^+^ARG1^+^) TAMs as a proportion of total TAMs in pups and adult mice bearing M3-9-M RMS. Data in C-I are presented as mean ± SEM. Sample sizes: n = 3-4 per group (A), n=12 per group *(B-ALL patients)*, n=4 younger, n=8 older *(AML patients)* (B), or n ≥ 5 per group (C-I). Statistical significance was determined using an unpaired Student’s t-tests (C-I). * p < 0.05, ** p < 0.01, *** p < 0.001, **** p < 0.0001.

### Host age influences the efficacy of anti-cancer immunotherapy

It has been widely observed that human tumors containing high levels of CD8^+^ T cells and inflammatory myeloid cells are more likely to respond to ICT^31,54^. Since we observed significant differences in the TIME between pediatric and adult mice, we hypothesized that host age would influence response to immunotherapy. To test this, we treated M3-9-M RMS-bearing mice of increasing age with an anti-PD-L1 antibody (**Fig. 4A**). Strikingly, while all adult mice exhibited complete tumor regression (**Fig. 4B**), no cures were achieved in pediatric mice inoculated with the same cell line (**Fig. 4C**). Moreover, we treated mice implanted at various developmental stages from early life to adulthood and observed an age-dependent response rate to ICT (**Fig. 4B-C, Fig. S6A**). Notably, this graded response was independent of PD-L1 expression on cancer cells, which was confirmed to be consistent across pups and adult mice (**Fig. S6B**). To determine whether tumor size or growth rate affected the age-dependent efficacy of ICT, adult mice were inoculated with ten-fold more cancer cells and treatment initiated once tumors were well established (∼150 mm^3^). Despite the larger tumor burden, ICT still induced complete regression in 60% of adult mice (**Fig. 4D, Fig. S6A**), indicating that whilst larger tumor burden affects response in adults, age has a significant influence on response to ICT. Similar results were observed in response to combination ICT (anti-CTLA-4 plus anti-PD-1) in two highly mutated sarcoma models, MCA205 and WEHI164, across C57BL/6 and BALB/c mouse strains, respectively (**Fig. 4E-F**).

**Figure 4.**
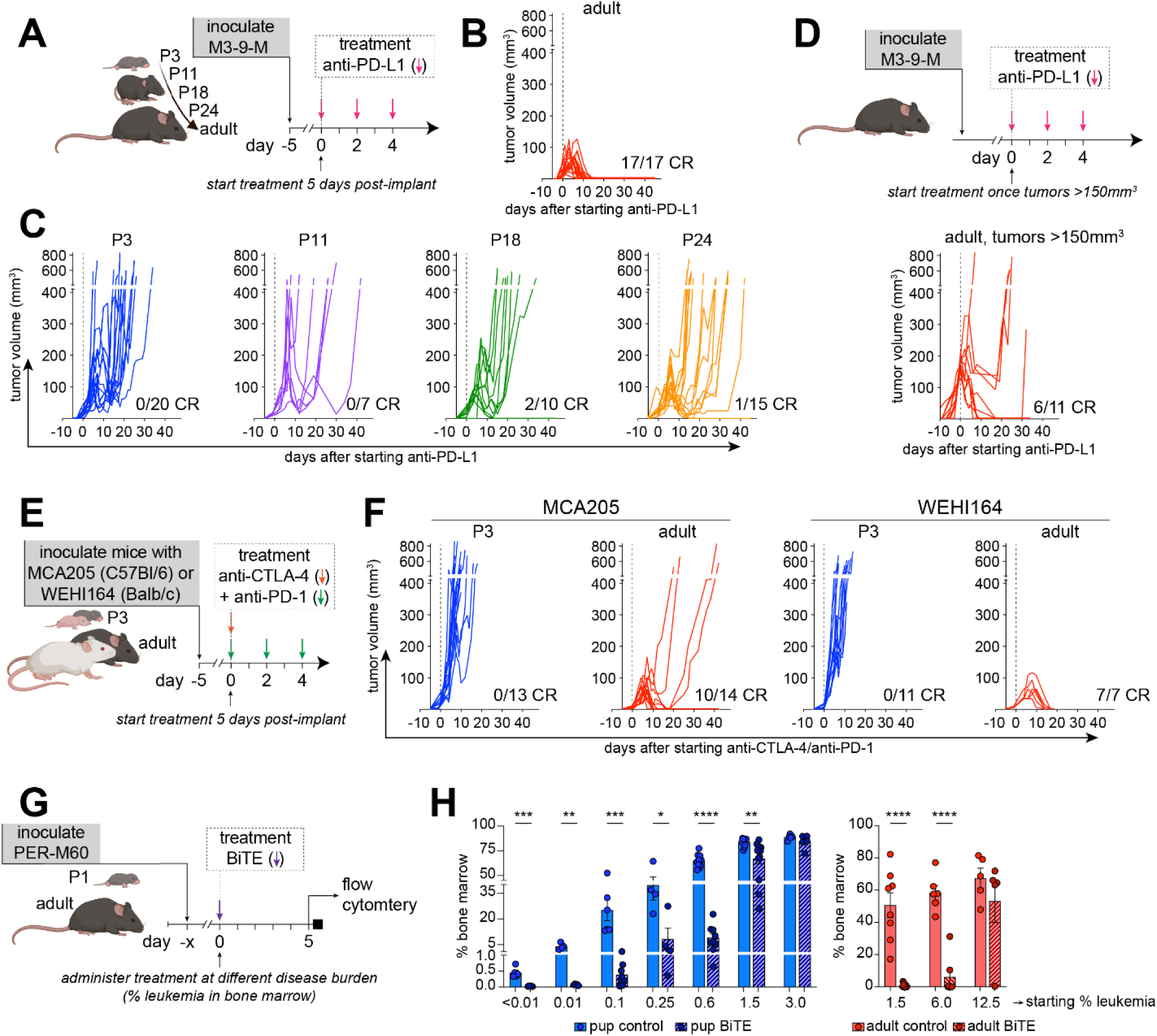
Anti-cancer immunotherapies are less effective in pediatric mice. (A) Experimental schema for B-C. M3-9-M RMS tumor volumes after anti-PD-L1 treatment (dotted line indicates start of treatment) in (B) adult mice and (C) young mice at different developmental stages (P3, P11, P18 and P24). Number of mice achieving complete response (*CR*) is indicated relative to total number assessed. (D) Experimental schema *(top)* and M3-9-M RMS tumor volumes *(bottom)* in adult mice treated with anti-PD-L1 with larger initial tumor burden. (E) Experimental schema for F. (F) Tumor volumes in mice implanted with the indicated cells and treated with combination ICT. (G) Experimental schema for H. (H) Percentage of PER-M60 B-ALL in the bone marrow after CD19×CD3 BiTE treatment, starting at different baseline leukemia burdens *(as indicated)*, in pediatric and adult mice. Data presented as mean ± SEM, n ≥ 7 per group (B, C, F), n ≥ 4 per group (H). Significance was calculated using an unpaired Student’s t-test. * p < 0.05, ** p < 0.01, *** p < 0.001, **** p < 0.0001.

To test whether these findings were consistent when using another class of immunotherapy relevant to pediatric cancers, we assessed a murine CD19×CD3 BiTE in PER-M60 B-ALL, designed to mimic the bispecific antibody blinatumomab, which has recently been established for use in the upfront setting for patients with B-ALL^5,55–58^. Pediatric and adult mice bearing PER-M60 B-ALL were treated with BiTE monotherapy (1 mg/kg) at several starting disease burdens in the bone marrow determined using flow cytometry (≤0.01-3% in pups and 1.5-12.5% in adult mice) (**Fig. 4G, Fig. S6C**). Pediatric mice exhibited an effective therapeutic response, but only when treatment was initiated at low leukemic burden; the response was effectively lost once the disease burden reached 1.5% at start of therapy. In contrast, adult mice showed a ∼99% reduction in leukemia, which only waned once the burden at start of treatment reached 12.5% (**Fig. 4H, Fig. S6D**). To exclude any age-dependent pharmacological effects, we treated pediatric mice with a five-fold higher dose, starting at 1% B-ALL disease burden in the bone marrow but no improvement in efficacy was observed (**Fig. S6E**). Blinatumomab has marked activity in pediatric B-ALL; however, in addition to confirming the known clinical association between tumor burden and blinatumomab response^59^, our results suggest its activity is negatively influenced by young age. To validate the translational relevance of our results, we analyzed response rates to blinatumomab in a unique cohort of patients with relapsed/refractory B-ALL who were treated with blinatumomab after induction chemotherapy as a bridge to hematopoietic stem cell transplantation, allowing assessment of the response specific to the blinatumomab^60^. This analysis showed that pediatric patients achieving a complete response after blinatumomab had a significantly higher age than patients who did not (**Fig. S6F**). Together, these data suggest that young host age is a critical determinant of reduced cancer immunotherapy efficacy.

### MYC shapes the TIME and response to ICT in pediatric tumors

Given these age-specific changes in the TIME across multiple pediatric cancers were associated with reduced response to immunotherapy, we sought to understand the molecular drivers of the pediatric TIME. We performed gene set enrichment analysis (GSEA)^61,62^ on bulk RNAseq data collected from tumors grown in pediatric and adult mice. These analyses revealed that gene sets indicative of MYC upregulation (MYC targets) and enhanced proliferation (G2M checkpoint and E2F targets) were highly enriched in tumors from pediatric mice (**Fig. 5A**). We further validated these findings using tumor gene expression data from pediatric cancer patients with embryonal and alveolar rhabdomyosarcomas, and medulloblastoma^63^, stratifying patients by age into 0-4 years (younger pediatric) and above 14 years (older pediatric) (**Table 1**). These cancer types were selected because they maintain similar histology and molecular features across the different age categories, and they are most representative of the murine models used in this study. Consistent with murine data, MYC targets and cell cycle-associated gene sets were enriched in younger children compared to older patients. Conversely, both adult mice and older patients displayed enrichment of gene sets indicative of immune activity and inflammatory response pathways, including gene sets associated with Interferon-α and Interferon-γ response among others (**Fig. 5A-B**).

**Figure 5.**
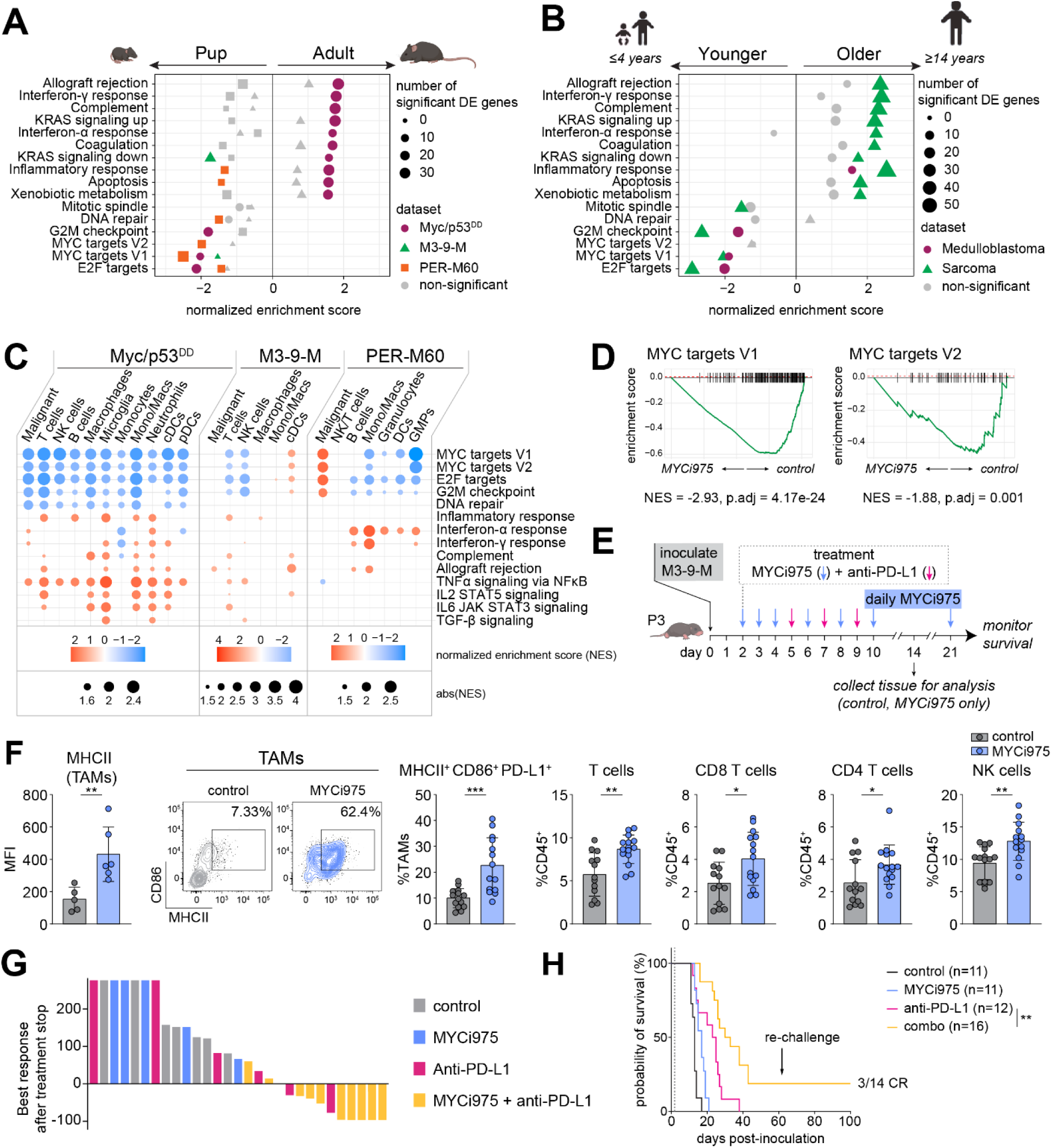
MYC activity is enhanced in the pediatric setting and inhibition modulates the TIME and ICT response. (A-B) GSEA of bulk RNAseq datasets from (A) mice and (B) pediatric patients of different age groups presented in order of the top 10 down- and top 6 up-regulated Hallmark gene sets in the murine Myc/p53^DD^ dataset. Symbols and colors represent cancer type as indicated. Significant enrichments (p_adj_ < 0.05) are indicated by colored symbols, with non-significant enrichments in grey. (C) GSEA of scRNAseq datasets from cancer-bearing pediatric and adult mice (blue: upregulated in pups, red: upregulated in adults), significantly different (p_adj_ < 0.05) gene sets associated with proliferation and immune response are presented. Sample sizes: n ≥ 5 /group (A); n = 79 (MB patients), n = 23 (RMS patients) (B); n ≥ 3 per group (C); n ≥ 6 per group. (D) GSEA of bulk RNAseq data from MYCi975-treated or control pediatric mice bearing M3-9-M RMS showing MYC targets V1 and V2. (E) Experimental schema for F-G (data are from at least 2 independent experiments). (F) The effect of MYCi975 (*pale blue*, compared to vehicle in *grey*) on MHCII abundance on TAMs, TAM phenotype *(left = representative plot, right = quantification)*, and other TILs (as indicated) in pediatric mice bearing M3-9-M RMS. Data are presented as mean ± SEM and significant differences determined using unpaired Student’s t-tests. (G-H) Pediatric mice bearing M3-9-M RMS were treated with vehicle control (*grey*), MYCi975 (*pale blue*), anti-PD-L1 (*pink*) or MYCi975 combined with anti-PD-L1 (combination, *yellow*). (G) Waterfall plot showing the best response for each condition after treatment cessation, calculated as percentage change in tumor volume and (H) survival curves are shown, with significant differences determined using log-rank (Mantel-Cox) test. * p < 0.05, ** p < 0.01, *** p < 0.001.

**Table 1:**
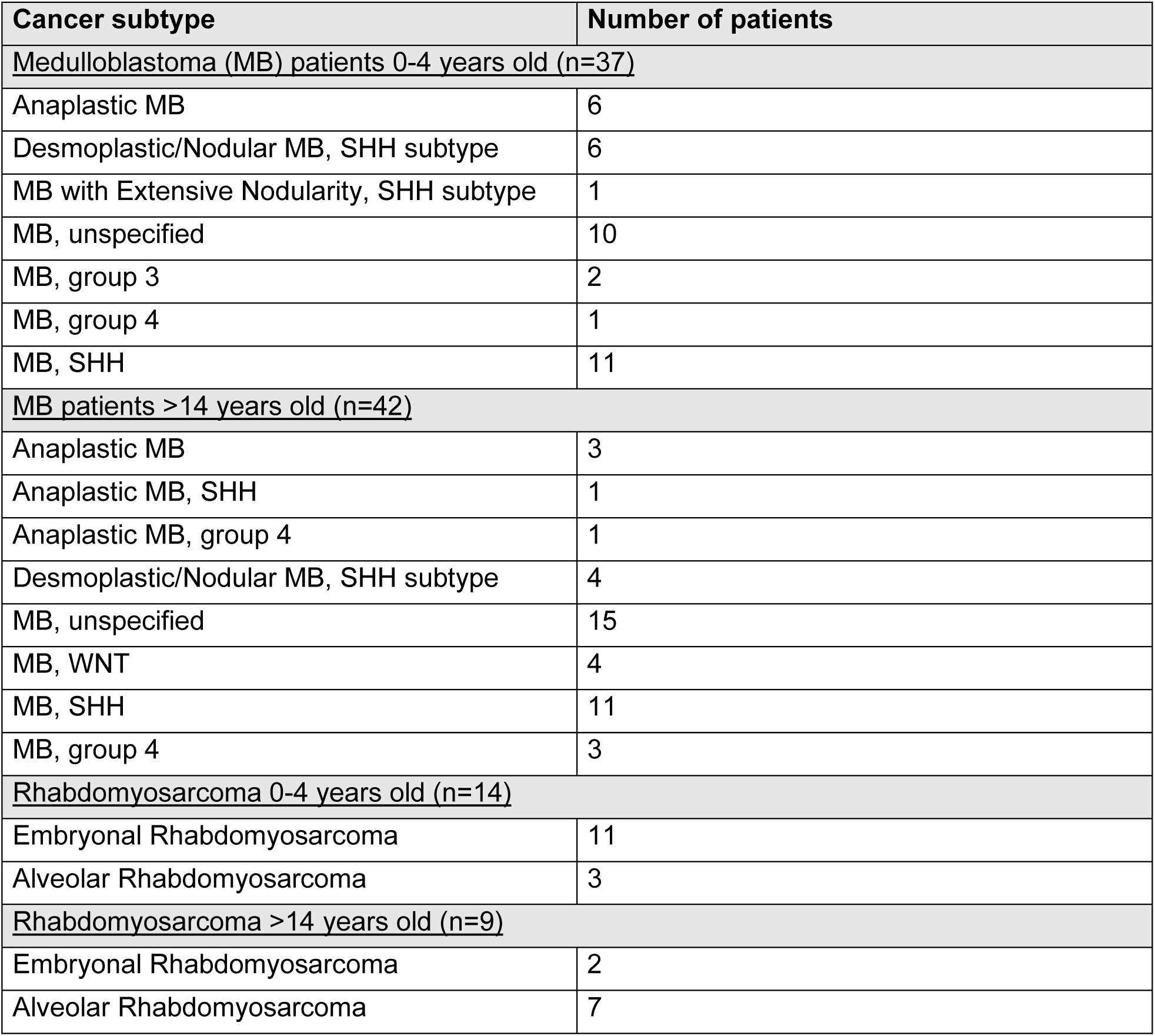
Patient characteristics. Diagnoses of the tumors used to generate Figure 5B as described in McLeod & Gout et al^63^.

Given the lack of tumor control in response to anti-PD-L1 we observed in pediatric mice compared to adult mice (**Fig. 4A-D**), we queried if these same gene sets were associated with treatment responsiveness in patients. Indeed, analysis of published bulk RNAseq data from pediatric patients treated with the anti-PD-L1 antibody atezolizumab revealed that MYC targets, E2F targets, and G2M gene sets were enriched in non-responsive tumors, while responsive tumors were enriched for inflammatory pathways (**Fig. S7A**), which was also shown by the study authors^64^. These murine and human data relied on bulk transcriptome analysis, preventing deeper understanding of which cells exhibited these gene expression changes. To identify the cell populations driving these gene expression differences, we conducted GSEA of our murine scRNAseq data and show that MYC target gene sets were particularly upregulated in TILs in pediatric mice. These gene sets were also enriched in other microenvironment cells, including stromal and endothelial cells. Moreover, MYC targets were enriched in G3MB Myc/p53^DD^ cells in pediatric mice, which was surprising given that *Myc* overexpression drives tumorigenicity in this model (**Fig. 5C**). Conversely, in adult mice, T cells, NK cells, dendritic cells, monocytes, and TAMs exhibited enrichment in inflammatory pathways (**Fig. 5C**). To validate these differences in patient samples, we performed single-sample GSEA (ssGSEA) using scRNAseq data from SHH MB patients comparing an infant with an older patient^49^. Consistent with results in mice, myeloid cells were enriched in MYC targets, E2F targets, and G2M gene sets in the infant SHH MB patient, while being deficient in inflammation-associated gene sets compared to the older patient (**Fig. S7B**). There were too few lymphocytes in these data, therefore we further validated the findings in a dataset of group 3/4 MB patients^65^. Again, MYC targets, E2F targets, and G2M gene sets were enriched in immune cells in younger patients (0-4 years old), compared to older patients (above 10 years old) (**Fig. S7C**). Furthermore, we analyzed single cell RNAseq datasets of leukemia samples from patients with B-ALL^47^ and AML^48^, comparing younger patients (0-4y) to older patients (above 14y). Again, immune pathways were enriched in immune cells from older B-ALL patients (≥14 years) (**Fig. S7D**), a finding that aligns with our murine B-ALL data, although the nature of the inflammatory signals varied, with TNF signaling and IL2-STAT5 signaling higher in older patients across both datasets, but IFN responses only elevated in the older B-ALL patients (**Fig. S7C-E**). Overall, these results underscore age-dependent differences in the tumor immune microenvironment, supporting our preclinical observations.

To better clarify the functional impact of MYC activity on tumor growth and TILs *in vivo*, we treated pediatric tumor-bearing mice with the potent and highly selective MYC inhibitor MYCi975^66–69^. Target inhibition was confirmed using bulk RNAseq of M3-9-M RMS after treating pediatric mice, showing downregulation of MYC target genes in treated mice (**Fig. 5D, Fig. S8A**). Additionally, targeting MYC in multiple solid tumor models (M3-9-M RMS and MCA205 fibrosarcoma) prolonged survival in immune-competent pediatric mice but not in an immune-deficient setting, suggesting the drug acts predominantly via immune cells (**Fig. S8B-D**). Although MYC inhibition did not affect the abundance of total TILs or TAMs (**Fig. S8E**), it did result in a notable increase in the frequency of intratumoral T cells (both CD4^+^ and CD8^+^) and NK cells (**Fig. 5F)**. In addition, treatment with MYCi975 induced a significant shift in TAM phenotype, marked by elevated MHCII expression and the emergence of a MHCII^+^/CD86^+^/PD-L1^+^ TAM subset, suggesting a transition towards a more pro-inflammatory state (**Fig. 5F**). These findings indicate that elevated MYC activity in pediatric mice contributes, in part, to the immunosuppressive TIME.

Given MYC activity appeared to modulate the pediatric TIME, and enrichment of MYC targets correlated with reduced responsiveness to ICT in pediatric mice and children (**Fig.S7A**)^64^, we hypothesized that MYC inhibition could increase ICT efficacy in pediatric cancer. We therefore implanted pediatric mice with M3-9-M RMS and administered anti-PD-L1 alone or in combination with MYCi975 **(Fig. 5E).** Combination therapy significantly improved disease control with 80% of mice exhibiting tumor regression (**Fig. 5G, Fig. S8F**). Moreover, 20% of mice demonstrated complete tumor rejection with sustained disease control, even after re-implantation with the same cancer cells (**Fig. 5H, Fig. S8F**). However, marked toxicity was seen in pediatric mice treated with the MYC inhibitor, with significantly stunted growth and failure to thrive (**Fig. S8G**), highlighting the central role that MYC plays in mediating tissue growth and development in early life^70^. Taken together, these data demonstrate that the high level of MYC activity associated with early life contributes to an immunosuppressive TIME, mediating unresponsiveness to ICT, which can be therapeutically reverted, although systemic MYC inhibition has prohibitive developmental impacts.

### TAM polarization overcomes the immunosuppressive pediatric TIME

Given the toxicity observed after pharmacological MYC inhibition, we sought alternative therapeutic avenues to increase the immunogenicity of the pediatric TIME. Since our results revealed a reduced inflammatory signature in pediatric cancers together with increased abundance of anti-inflammatory MHCII^low^/M2-like TAMs, we hypothesized that polarizing TAMs towards a pro-inflammatory MHCII^hi^ phenotype would increase CD8^+^ T cell infiltration and cytotoxicity. Multiple studies have identified that treatment with an agonistic antibody against CD40 in combination with a blocking antibody against colony stimulating factor 1 receptor (CSF1R) achieves this pro-inflammatory TAM polarization^71–73^. We therefore treated pediatric mice harboring Myc/p53^DD^ G3MB, M3-9-M RMS, and PER-M60 B-ALL with anti-CD40 and anti-CSF1R antibodies. Treatment significantly decreased disease burden in pediatric mice with M3-9-M RMS and PER-M60 B-ALL, and extended survival in mice with Myc/p53^DD^ G3MB (**Fig. 6A-B, Fig. S9A-B**). Importantly, anti-CD40/anti-CSF1R treatment increased MHCII expression on TAMs, in addition to the levels of CD40, PD-L1, PD-L2, CD86, and CD80 in all three cancer types (**Fig. 6C, Fig S9C-E**). Notably, the proliferative (Ki67^+^) MHCII^low^ TAMs were the most sensitive population to be depleted by anti-CD40/anti-CSF1R (**Fig. 6D**), consistent with a previous report^74^. These changes in TAMs induced by anti-CD40/anti-CSF1R indicate a potential shift towards an M1-like phenotype, consistent with our hypothesis.

**Figure 6.**
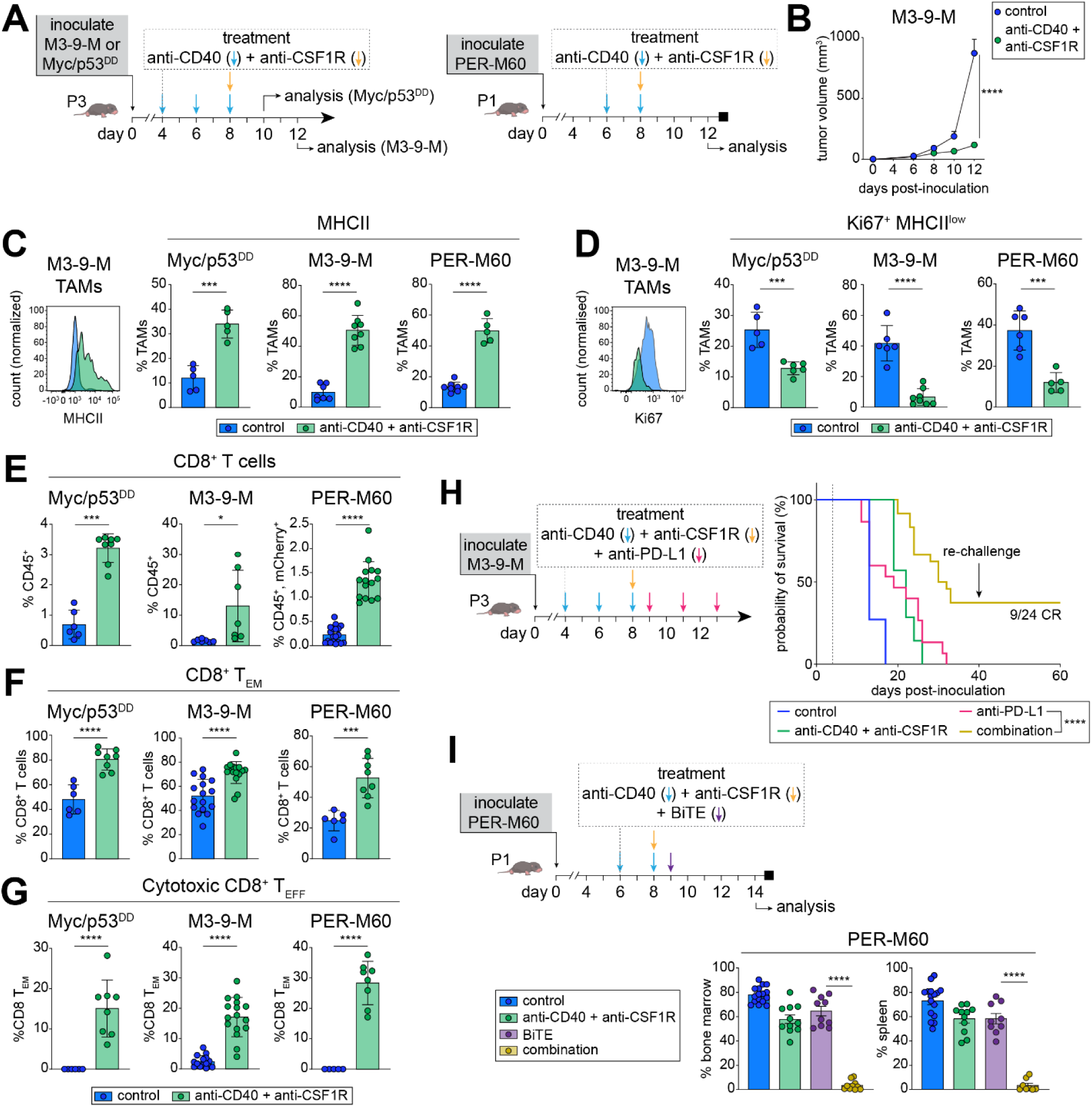
TAM polarization overcomes the immunosuppressive pediatric TIME. (A) Experimental schema for (B-G). (B) Tumor growth curve of M3-9-M RMS after anti-CD40/anti-CSF1R treatment in pediatric mice. (C) Representative flow cytometry histograms of MHCII expression on TAMs (*left*, *M3-9-M* shown) and frequency of MHCII^+^ TAMs as a proportion of total TAMs in the murine models indicated following treatment (*green*) compared to vehicle controls (*blue*). (D) Representative flow cytometry histogram of Ki67 expression in TAMs (*left, M3-9-M shown*) and frequency of Ki67^+^ MHCII^low^ TAMs as a proportion of total TAMs in the indicated murine models following treatment. (E-G) Frequency of (E) CD8^+^ T cells as a percentage of CD45^+^ cells, (F) T_EM_ as a percentage of total CD8^+^ T cells, and (G) T_EFF_ (KLRG1^+^Ki67^+^GZMA^+^) as a percentage of total CD8^+^ T_EM_ cells in pups bearing Myc/p53^DD^ G3MB, M3-9-M RMS, or as a percentage of CD45^+^mCherry^-^ cells in pups bearing PER-M60 B-ALL following treatment. (H) Schema (*left*) and survival curves (*right*) of pediatric mice bearing M3-9-M RMS after treatment with vehicle (*blue*), anti-CD40/anti-CSF1R (*green*), anti-PD-L1 (*pink*) or the combination (*yellow*). (I) Schema *(top)* and percentage of leukemia cells in the bone marrow and spleen (*bottom)* of pediatric mice bearing PER-M60 B-ALL after treatment with vehicle (*blue*), BiTE (*plum*), anti-CD40/anti-CSF1R (*green*), or the combination (*yellow*). Data are presented as mean ± SEM. Sample sizes: n ≥ 5 per group (B–G), n ≥ 8 per group (H and I). Statistical significance was calculated using mixed-effects models (B), unpaired Student’s t-tests (C–G and I), and a log-rank (Mantel-Cox) test (H). * p < 0.05, ** p < 0.01, *** p < 0.001, and **** p < 0.0001.

Critically, M1-like polarization was associated with increased CD8^+^ T cell abundance across all three types of cancer (**Fig. 6E**). More specifically, treatment appeared to promote a T_EM_ phenotype, with an emergence of highly proliferative cytotoxic KLRG1^+^ effector population (cytotoxic T_EFF_: CD44^+^/CD62L^-^/KLRG1^+^/Ki67^+^/GZMA^+^) (**Fig. 6F-G**). Concordantly, the treatment reduced T_EX_, but did not lead to T_RM_ expansion (**Fig. S9F**). Next, we assessed how this therapeutic polarization of the pediatric TIME impacted response to anti-PDL1 in M3-9-M RMS, and BiTE in B-ALL. Pediatric mice with M3-9-M RMS were treated with anti-CD40/anti-CSF1R and anti-PD-L1. While no complete responses were observed with anti-PD-L1 or anti-CD40/anti-CSF1R treatments alone, 40% of pediatric mice treated with the triple combination achieved complete tumor rejection and developed immune memory, as evidenced by their rejection of re-implanted M3-9-M RMS cells on the contralateral flank (**Fig. 6H**). The treatment was well tolerated by pediatric mice. To test whether macrophage polarization could sensitize pediatric tumors to other immunotherapies, pediatric mice with PER-M60 B-ALL were treated with anti-CD40/anti-CSF1R followed by CD19×CD3 BiTE (**Fig 6I**). Whilst anti-CD40/anti-CSF1R or BiTE did not have a marked effect on leukemia progression, the combination displayed a 95% reduction of leukemia cells in the bone marrow and spleen compared to control (**Fig. 6I**). The efficacy of the combination was also confirmed in a second model of B-ALL (PER-M224^66^, **Fig. S9G**). Overall, our data show that targeting young age-dependent immune suppression, mediated in part by TAMs, can modulate the TIME, and most importantly, this can improve efficacy of clinically available immunotherapies across various pediatric cancers.

## DISCUSSION

In this study, we have established a suite of pediatric mouse cancer models specifically designed to reflect age-related changes in pediatric cancer immunobiology. Importantly, we provide evidence for age-dependent effects shaping the TIME with relevant therapeutic implications.

Analysis of pediatric tumors from young patients compared with older age groups (in RMS^75^) or with adult patients (in neuroblastoma^44^, Wilms tumor^44^, diffuse intrinsic pontine glioma^45^, and B-ALL^76^) has indicated age-dependent differences in TILs. Pediatric tumors are typically heavily infiltrated by macrophages^44,45,76^, while having fewer T cells^44,45,75^ and dendritic cells^44^. A recent study identified age-dependent differences in circulating immune cells in children with cancer, with for example lower CD8, CD4 T cells and NK cell numbers compared to healthy controls^77^. These differences raise the question of whether the distinct TIME observed in pediatric cancers is driven by the host’s age or by intrinsic cancer biology. To address this, we used inbred mouse models of varying ages, allowing us to isolate age as an experimental variable, while keeping other variables consistent, including cancer cell and germline genetics. Across multiple pediatric cancer types, our models consistently demonstrated significant age-related differences in the TIME, consistent with clinical observations seen in children with cancer^44,45,78^.

Previous SARS-CoV-2 studies have highlighted age-dependent differences in systemic T cell responses, showing delayed CD8^+^ T cell effector and memory formation accompanied by enrichment of naïve CD8^+^ T cells in young individuals^26,79^. The dynamic changes in tissue T cell populations and phenotypes during early life^27,28^, coupled with the relative rarity of pediatric cancers throughout childhood, and generally low immune cell infiltration in these tumors^46,78^, have posed significant challenges for conducting high-dimensional analyses of pediatric TILs. In our pediatric cancer models, tumor-infiltrating CD8^+^ T cells predominantly exhibit a naïve-like phenotype characterized by limited expression of immune checkpoints such as PD-1 and LAG3. This was accompanied by less effector and exhausted CD8^+^ T cells, and a reduced formation of specialized T cells such as T_RM_. Robust anti-tumor T cell responses following ICT and adoptive T cell therapies are dependent on CD8^+^ T cells that retain proliferative and differentiation capacity^31^. Indeed, anti-CD40/anti-CSF1R increased effector and cytotoxic CD8^+^ T cells, suggesting that naïve like CD8^+^ TILs in young mice retain the capacity to differentiate into more specialized populations, such as effector memory T cells.

Our study also uncovers a distinct profile of myeloid cells in pediatric mouse tumors, which was corroborated in B-ALL, AML, and SHH MB patient datasets, marked by notably low expression levels of MHCII. This finding aligns with previous clinical studies demonstrating that cord blood monocytes and alveolar macrophages in infants exhibit reduced HLA-DR expression, leading to decreased antigen presentation capabilities and a consequently less robust adaptive immune response^80,81^. Moreover, in children with B-ALL, bone marrow monocytes have been shown to exhibit low expression of MHCII genes (HLA-DQA1 and HLA-DPB1), which was associated with enriched naïve T cells^82^. Interestingly, MHCII^low^ macrophages have also been identified within immune-suppressive, M2-like TAMs, in both adult murine cancer models and adult human patients^83–85^. However, our data suggests that TAMs in early life are predisposed toward an M2-like phenotype, inherently exhibiting low MHCII expression, which may contribute to the lower adaptive immune response observed in pediatric cancers. Considering the prenatal origin of many pediatric cancers^86^, additional studies using cell fate mapping in mouse models are required to trace how embryonically derived macrophages shape the TIME in pediatric cancers. Understanding these mechanisms could provide critical insights into tailoring immunotherapeutic strategies for young patients.

Single cell transcriptomic analysis across multiple murine models and tumors of young cancer patients identified the MYC pathway as the most consistently upregulated biological pathway in the pediatric setting, and we pinpointed this upregulation predominantly to TILs. MYC is not only an oncoprotein that drives cancer cell proliferation, but also a key regulator of embryogenesis and early-life development, orchestrating immune cell death, proliferation, and metabolism^70^. For example, knockout studies revealed that loss of *Myc* promotes the emergence of pro-inflammatory, M1-like macrophages^87–89^. The increased MYC activity observed in tumors in our pediatric mouse models and in young cancer patients is associated with an upregulation of E2F and G2M cell cycle pathways, with lower inflammatory responses compared to older mice and older patients. This suggests that MYC orchestrates a fine balance between growth and immune suppression^87^ which is crucial for normal development but could also be contributing to the unique immune environment observed in pediatric cancers. The dual role of MYC in promoting both cellular growth and immune suppression presents significant challenges for targeting it in pediatric cancer. Although pharmacological MYC inhibition enhanced the efficacy of anti-PD-L1 immunotherapy in our pediatric mouse models, it also resulted in considerable toxicity. This highlights the complexity of targeting MYC in pediatric cancers, where its role in normal development must be carefully weighed against its contribution to tumorigenesis and immune evasion^90^.

As an alternative, we sought to modulate the pediatric TIME with antibodies targeting CD40 and CSF1R. These treatments proved highly effective in polarizing macrophages towards a pro-inflammatory phenotype, and importantly, this modulation sensitized tumors to clinically appropriate immunotherapy with either ICT or BiTE. Although rigorous toxicology assessments will be necessary, the fact that both CD40 and CSF1R antibodies are already in clinical trials^91,92^, including in pediatric studies^93^, provides a strong rationale for clinical translation.

Given the limited efficacy of ICT in pediatric cancer, except in rare cases of tumors with constitutional mismatch repair deficiency^94^ and classic Hodgkin lymphoma^95^, there is an urgent need to better harness the immune response in children with cancer. Even though children with B-ALL can be effectively treated with blinatumomab in combination with chemotherapy^5,58,96,97^, response to blinatumomab is not necessarily universal. This is highlighted by the first early phase pediatric trial of blinatumomab monotherapy in relapsed/refractory B-ALL, where there was a 39% complete response rate^98^, which is lower than observed in adults^99^. Considering that blinatumomab will now form a standard component of upfront therapy for children with B-ALL, it is imperative to identify features that may help to predict and improve response.

Unlike adult cancers, where thousands of clinical trials are exploring a multitude of treatment combinations, pediatric cancer trials are fewer in number due to the rarity of these diseases^100,101^. This scarcity of trials, coupled with the slower pace of patient recruitment, makes it inefficient to clinically test numerous treatment combinations. Consequently, the field relies heavily on predictive and translationally relevant preclinical models to guide clinical trial design, helping to prioritize the most promising treatment regimens while minimizing the risk of toxicities. By leveraging key insights from developmental immunology and infectious diseases^26–28,79,102–105^, along with the use of age-appropriate pediatric mouse models that better reflect the dynamic immune development in children, we propose that we can accelerate the creation of tailored immunotherapeutic strategies with higher chances of clinical success. This approach could pave the way for more effective treatments that offer long-term cures without the lifelong side effects associated with current standard-of-care therapies like chemotherapy or radiotherapy.

### Limitations of the Study

Our study utilized pediatric cancer models to explore age-specific effects on tumor immunobiology. However, the complex interplay between host-derived factors—such as growth factors, cytokines, and chemokines, and cancer cell-intrinsic oncogenic signaling perturbations^106^ suggests these models could also be employed to comprehensively characterize non-immune interactions in future studies. We chose transplantable models to reduce variability in cancer progression and mutations, since genetically-engineered mouse models of pediatric cancer often develop tumors at varying (often adult) ages and are typically associated with the spontaneous acquisition of additional mutations that differ from mouse to mouse^107^. While our focus was primarily on CD8^+^ T cells and macrophages, we also observed age-dependent differences in other tumor-infiltrating immune cell populations, which warrant further characterization and functional validation in subsequent studies. Finally, although we demonstrated the impact of age on the TIME, these mouse models could also be used to explore how early-life biology influences cancer development, providing a broader understanding of the relationship between age and tumorigenesis.

## ACKNOWLEDGEMENTS

The authors would like to acknowledge the generous support of the Stan Perron Charitable Foundation, the Australian Lions Childhood Cancer Research Foundation, The Pirate Ship Foundation, Telethon, The Kids Cancer Project, the Child Cancer Research Foundation, Kerimi Family Foundation, the Western Australian Future Health Research and Innovation Fund (FHRI), the Western Australian Child Research Fund, Cancer Council Western Australia and the Genome Canada Science Technology Innovation Centre, Compute Canada Resource Allocation Project. This work was also supported by resources provided by the Pawsey Supercomputing Research Centre’s Nimbus Research Cloud (doi:10.48569/v0j3-qd51), with funding from the Australian Government and the Government of Western Australia through allocation to M.A.W. W.J.L. is supported by a National Health and Medical Research Council (NHMRC) Investigator Grant (2021/GNT1196605). M.R.J. has received funding from NHMRC Investigator Grant APP1172858 and Synergy Grant APP2010849. T.L. and N.G.G. are supported by the Stan Perron Charitable Foundation. R.E. is supported by a Cancer Council WA Research Fellowship and a Pirate Ship Foundation Brainchild Fellowship. A.S. is supported by a Pirate Ship Fellowship. N.G.G. is supported by Perth Children’s Hospital Foundation Stan Perron Chair in Paediatric Oncology and Haematology. R.S.K. is supported by an NHMRC Investigator Grant APP2033152. C.L.K has received funding from the Canadian Institutes of Health Research (CIHR) grant PJT-190271. ZA was supported by a Richard Walter Gibbon Medical Research Scholarship and an Australian Government Research Training Program Scholarship at the University of Western Australia. This research was made possible, in part, through the financial support provided by the Brain Cancer Centre, facilitated by members M.R.J., N.G.G, and R.E. The funders had no role in the study design, data collection, analysis, decision to publish, or manuscript preparation.

Library preparation and sequencing was conducted in the Genomics WA Laboratory in Perth, Australia. This facility is supported by BioPlatforms Australia, State Government Western Australia, Advanced Cancer Research Foundation, Cancer Research Trust, Harry Perkins Institute of Medical Research, The Kids Research Institute Australia and the University of Western Australia. We gratefully acknowledge the Australian Cancer Research Foundation and the Centre for Advanced Cancer Genomics for making available Illumina Sequencers for the use of Genomics WA.

We acknowledge the significant contribution made by all the animals used in this study. Thank you to the Bioresources teams of The Kids Research Institute Australia for the care of our animals, animal welfare officer Deborah McDonald, senior program manager Emma Stone and the Laboratory Management team, particularly Pradeep Kumar, for maintenance of the flow cytometry facility. We thank all members of the WA Kids Cancer Centre, Clara Andradas Arias, Imran Khan, Brittany Dewdney, Linda Wijaya, Ankur Sharma and Rhea Pai for their advice, suggestions, and discussions during this project.

We also wish to express gratitude to the patients and their families, the medical staff and the WA Kids Cancer Community Reference Group.

## AUTHOR CONTRIBUTIONS

Conceptualization, O.E., Z.A., T.J., R.E., T.L., W.J.L., data curation, O.E., Z.A., M.A.W.; formal analysis, O.E., Z.A., M.A.W., A.S., T.J., R.S.K., L.C.C., S.M., T.L., R.E., W.J.L.; funding acquisition, O.E., M.H., M.R.J., N.G.G., T.J., R.S.K., L.C.C., S.M., R.E., T.L., W.J.L., investigation, O.E., Z.A., M.A.W., J.T., I.M.J., J.K., A.W., H.S., H.H., I.P. G.A.C., V.N., J.O., I.K., K.P.; methodology, O.E., Z.A., M.A.W, J.T., I.M.J., J.K., H.S., H.H., S.S., F.A., A.M., A.S., R.M.Z., B.W., J.C., B.F., T.L., L.C.C., S.M., T.L., R.E., W.J.L., project administration, O.E., R.E., T.L., W.J.L., software, O.E., Z.A., M.A.W., A.N., C.L.K.; supervision, O.E., S.M., R.S.K., L.C.C, R.E., T.L., W.J.L.; validation, O.E., Z.A., M.A.W., J.T., I.M.J., J.K., A.N., C.L.K., N.J.; visualization, O.E., Z.A., R.E., W.J.L.; resources, O.E., T.N.P., L.C.C., S.M., R.E., W.J.L., writing, O.E., Z.A., R.S.K., S.M., T.L., R.E., W.J.L.; review and editing, all.

## DATA AVAILABILITY

The data generated and analyzed in this study (Bulk RNAseq and scRNAseq data from mouse tumors) will be available upon request. Feature count data for diagnostic rhabdomyosarcoma and medulloblastoma tumor samples used for analysis in this study were obtained from St. Jude Cloud^63^ (https://www.stjude.cloud). This study makes use of data generated by the St. Jude Children’s Research Hospital – Washington University Pediatric Cancer Genome Project, Childhood Solid Tumor Network and/or Real-time Clinical Genomics^108–110^. RNA sequencing gene expression data from atezolizumab (non-) responders was obtained via the European Genome-Phenome Archive, accession number EGAS00001006004. scRNAseq data from B-ALL and AML patients is available through the Alex’s Lemond Stand Single-cell Pediatric Cancer Alex’s Lemond Stand Single-cell Pediatric Cancer ^111^, projects SCPCP00000^47,48^, scRNAseq data from G3/G4 MB patients in the GEO repository via the accession number GSE155446, the GEO repository via the accession number GSE155446.

## STAR METHODS

### Key Resource table

| REAGENT or<br>RESOURCE | SOURCE | IDENTIFIER |
| --- | --- | --- |
| Flow cytometry antibodies |  |  |
| Anti-mouse CD45-BUV395 (clone 30-F11) | BD Biosciences | Cat: 564279 RRID: AB_2651134 |
| Anti-mouse CD45-PE (clone 30-F11) | BD Biosciences | Cat: 553081 RRID: AB_394611 |
| Anti-mouse CD45-BV570 (clone 30-F11) | BioLegend | Cat: 103136 RRID: AB_2562612 |
| Anti-mouse<br>CD45.1- BV650<br>(clone A20) | BD Biosciences | Cat: 563754 RRID: AB_2738405 |
| Anti-mouse<br>CD45.2- Alex 700<br>(clone 104) | BD Biosciences | Cat: 560693 RRID: AB_1727491 |
| Anti-mouse CD11b-BUV496 (clone<br>M1/70) | BD Biosciences | Cat: 749864 RRID: AB_2874105 |
| Anti-mouse CD11b-BV510 (clone M1/70) | BD Biosciences | Cat: 562950 RRID: AB_2737913 |
| Anti-mouse CD11b-RB744 (clone M1/70) | BD Biosciences | Cat: 570512, RRID:AB_3665015 |
| Anti-mouse CD11b-BUV395 (clone M1/70) | BD Biosciences | Cat: 569276, RRID: AB_3626323 |
| Anti-mouse CD93-BUV661 (clone AA4.1) | BD Biosciences | Cat: 741574 RRID: AB_2870995 |
| Anti-mouse CRig-BUV737 (clone 17C9) | BD Biosciences | Cat: 749499 RRID: AB_2873867 |
| Anti-mouse F4/80-BUV805 (clone T45-2342) | BD Biosciences | Cat: 749282 RRID: AB_2873657 |
| Anti-mouse F4/80-BB700 (clone T45-2342) | BD Biosciences | Cat: 746070 RRID: AB_2743450 |
| Anti-mouse TIM-4-BUV805 (clone 21H12) | BD Biosciences | Cat: 748750 RRID: AB_2873154 |
| Anti-mouse Ly6C-BV421 (clone AL-21) | BD Biosciences | Cat: 562727 RRID: AB_2737748 |
| Anti-mouse Ly6C-RY775 (clone AL-21) | BD Biosciences | Cat: 571412, RRID:AB_3665016 |
| Anti-mouse Ly6C-RY703 (clone AL-21) | BD Biosciences | Cat: 571456 |
| Anti-mouse Ly6C-FITC (clone HK1.4) | BioLegend | Cat: 128006 RRID: AB_1186135 |
| Anti-mouse MHCII-BV510 (clone M5/114.15.2) | BD Biosciences | Cat: 742893 RRID: AB_2741133 |
| Anti-mouse MHCII-BV605 (clone M5/114.15.2) | BD Biosciences | Cat: 563413 |
| Anti-mouse CD11C- BV605 (clone N418) | BioLegend | Cat: 563413 RRID: AB_2738190 |
| Anti-mouse PD-L1-RB780 (clone<br>MIH5) | BD Biosciences | Cat: 755394 |
| Anti-mouse CD86-AF647 (clone GL-1) | BioLegend | Cat: 105020, RRID: AB_493464 |
| Anti-mouse CD86-APC (clone GL-1) | Cell Signaling<br>Technology | Cat: 84393, RRID: AB_3665017 |
| Anti-mouse Ly-6G-AF700 (clone 1A8) | BD Biosciences | Cat: 561236, RRID: AB_10611860 |
| Anti-mouse<br>Arginase 1- eF450<br>(clone A1exF5) | Thermo Fisher | Cat: 48-3697-82, RRID: AB_2734837 |
| Anti-mouse CD206-BB630-P2 (clone<br>Y17-505) | BD Biosciences | Custom BB630-P2 Conjugate |
| Anti-mouse FR-β-<br>PE (clone 10/FR2) | BioLegend | Cat: 153304, RRID: AB_2721344 |
| Anti-mouse NK1.1-PE-Cy7 (clone<br>PK136) | BD Biosciences | Cat: 552878, RRID: AB_394507 |
| Anti-mouse PD-1-BUV395 (clone<br>29F.1A12) | BD Biosciences | Cat: 568595 |
| Anti-mouse CD4-BUV496 (clone<br>GK1.5) | BD Biosciences | Cat: 612952, RRID: AB_2813886 |
| Anti-mouse CD44-BUV563 (clone<br>IM7) | BD Biosciences | Cat: 569705, RRID: AB_3626317 |
| Anti-mouse LAG-3<br>- BUV661 (clone<br>C9B7W) | BD Biosciences | Cat: 741594, RRID: AB_2871003 |
| Anti-mouse CD3-BUV737(clone<br>17A2) | BD Biosciences | Cat: 612803, RRID: AB_2738781 |
| Anti-mouse CD8a-BV480 (clone 53-6.7) | BD Biosciences | Cat: 566096, RRID: AB_2739500 |
| Anti-mouse CD62L-BV605 (clone MEL-14) | BD Biosciences | Cat: 563252, RRID: AB_2738098 |
| Anti-mouse CD49a-BV650 (clone<br>Ha31/8) | BD Biosciences | Cat: 740519, RRID: AB_2740235 |
| Anti-mouse TIM3-BV650 (clone<br>5D12/TIM-3) | BD Biosciences | Cat: 747626 |
| Anti-mouse<br>KLRG1- BV711<br>(clone 2F1) | BD Biosciences | Cat: 564014, RRID: AB_2738542 |
| Anti-mouse CD69-RB780 (clone<br>H1.2F3) | BD Biosciences | Cat: 755339, RRID: AB_3626319 |
| Anti-mouse<br>CXCR6- PE (clone<br>SA051D1) | BioLegend | Cat: 151104, RRID: AB_2566546 |
| Anti-mouse NK1.1-PE-Cy7 (PK136) | BD Biosciences | Cat: 552878, RRID: AB_394507 |
| Anti-mouse<br>RUNX3- BV421<br>(clone R3-5G4) | BD Biosciences | Cat: 565742, RRID: AB_2916369 |
| Anti-mouse Ki-67-RB545 (clone B56) | BD Biosciences | Cat: 569276, RRID: AB_3626323 |
| Anti-mouse GZMA-PerCP eF710<br>(clone GzA-3G8.5) | Thermo Fisher | Cat: 46-583182, RRID: AB_2573775 |
| Anti-mouse TOX-eF660 (clone TXRX10) | Thermo Fisher | Cat: 50-6502-82, RRID: AB_2574265 |
| Fixable Viability Dye eF780 | Thermo Fisher | Cat: 65-0865-14 |
| Critical commercial kits |  |  |
| RNeasy Micro Kit | Qiagen | Cat: 74004 |
| RNeasy Mini Kit | Qiagen | Cat: 74104 |
| Foxp3/Transcription Factor Staining Buffer Set | BD Biosciences | Cat: 560409, RRID: AB_2869340 |
| MojoSort™ Mouse CD3 T Cell Isolation Kit (Biolegend). | Biolegend | Cat: 480031 |
| Experimental models: Cell lines |  |  |
| M3-9-M RMS | Kindly shared by Crystal Mackall<sup>22</sup>, Stanford University | NA |
| MCA205 fibrosarcoma | Sigma-Aldrich | NA |
| WEHI164 fibrosarcoma | CellBank Australia | NA |
| PER-M60 B-ALL | Kindly provided by<br>Laurence<br>Cheung<sup>20</sup>, The Kids<br>Research Institute<br>Australia | NA |
| PER-M224 B-ALL | Kindly provided by<br>Sebastien<br>Malinge<sup>112</sup>, The<br>Kids Research<br>Institute Australia | NA |
| PER-M5 AML | Kindly provided by<br>Sebastien Malinge,<br>The Kids Research<br>Institute Australia | NA |
| Myc/p53<sup>DD</sup> G3MB | Kindly provided by<br>Robert Wechsler-Reya, Columbia<br>University, USA | NA |
| H3.1<sup>K27M</sup>DMG | Kindly provided by<br>Timothy N<br>Phoenix<sup>24</sup>,<br>Cincinnati<br>Children's Hospital | NA |
|  | Medical Center,<br>Cincinnati, Ohio,<br>USA |  |
| TH-MYCN 9464D<br>neuroblastoma | Kindly provided by<br>Paul Sondel<sup>25</sup>,<br>Department of<br>Pediatrics,<br>University of<br>Wisconsin-Madison, Madison,<br>Wisconsin, USA | NA |
| Experimental models: Organisms/strains |  |  |
| Mouse: BALB/cJ | The Jackson<br>Laboratory and<br>maintained at The<br>Kids Research<br>Institute Australia,<br>bioresources facility | Strain# 000651 RRID: IMSR_JAX: 000651 |
| Mouse: C57BL/6J | The Jackson<br>Laboratory and<br>maintained at The<br>Kids Research | Strain# 000664 RRID: IMSR_JAX:000664 |
|  | Institute Australia,<br>bioresources facility |  |
| Mouse: B6 albino | The Jackson<br>Laboratory and<br>maintained at The<br>Kids Research<br>Institute Australia,<br>bioresources facility | Strain# 000058 RRID: IMSR_JAX: 000058 |
| Mouse: B6 CD45.1 | Ozgene ARC | Strain# 002014 RRID: IMSR_JAX: 002014 |
| Mouse: NOD/LtSz-Rag1<br>null (NOD/Rag1KO) | The Jackson<br>Laboratory and<br>maintained at The<br>Kids Research<br>Institute Australia,<br>bioresources facility | Strain# 003729 RRID: IMSR_JAX:003729 |
| Software and algorithms (see text for individual packages) |  |  |
| FACSDiva 6 | BD Biosciences | RRID:SCR_001456 |
| Cell Ranger 6.1.1 | 10X Genomics | RRID:SCR_017344 |
| Other |  |  |
| Neuros Syringe 2<br>µL, Model 75 RN,<br>33 gauge, point | Hamilton | Cat: 65460-03 |
| style 4 (for pediatric implantation) |  |  |
| 10 µL Microliter Syringe Model 701 N, Cemented Needle, 26s gauge, 2 in, point style 2 (for adult implantation) | Hamilton | Cat: 80300 |
| Ideal Micro-Drill™ hand-held microdrill | CellPoint Scientific | Cat: RS-6300-UK |
| Brilliant Stain Buffer Plus | BD Biosciences | Cat: 566385 |
| FC Block | BD Biosciences | Cat: 553141, RRID: AB_394656 |
| Collagenase Type IV | Sigma-Aldrich | Cat: C4-22-1G |
| DNase I Solution (1 mg/mL) | Stemcell Technologies | Cat: 07900 |
| DNase I (lyophilized) | Sigma-Aldrich | Cat: DN25-100MG |
| Hyaluronidase | Sigma-Aldrich | Cat: H3506-1G CAS: 37326-33-3 |
| Liberase | Roche | Cat: 05401020001 |
| Hank's buffered salt solution (HBSS) | Gibco | Cat: 14175095 |
| EDTA disodium salt | Sigma-Aldrich | Cat: E5134 |
| 10X RBC Lysis<br>Buffer | Thermo Fisher<br>Scientific | Cat: 00-4300-54 |
| MACS®<br>SmartStrainers (70<br>µm) | Miltenyi | Cat: 130-110-916 |
| Bovine Serum<br>Albumin | Bovogen<br>Biologicals | Cat: BSAS-1.0 |
| Sodium Azide,<br>Extra Pure | Thermo Fisher<br>Scientific | Cat: S-2360-48 CAS: 26628-22-8 |
| DPBS, powder, no<br>calcium, no<br>magnesium | Thermo Fisher<br>Scientific | Cat: 21600044 |
| TissueRuptor | Qiagen | Cat: 9002756 |
| gentleMACS Octo<br>Dissociator | Miltenyi Biotec | Cat: 130-096-427 |
| gentleMACS C<br>tubes | Miltenyi Biotec | Cat: 130-093-237 |
| RPMI 1640 Medium | Thermo Fisher<br>Scientific | Cat: 11875119 |
| Fetal Bovine Serum | CellSera | Cat: AU-FBS/PG |
| RBC lysis solution | Miltenyi | Cat: 130-094-183 |
| Trypsin-EDTA<br>(0.25%) | Thermo Fisher<br>Scientific | Cat: 25200072 |
| HEPES (1 M) | Thermo Fisher Scientific | Cat: 15630080 |
| MEM Non-Essential Amino Acid Solution (100X) | Sigma-Aldrich | Cat: M7145 |
| Neurobasal™ media | Gibco | Cat: 21103049 |
| Penicillin-Streptomycin (10,000 U/mL) | Thermo Fisher Scientific | Cat: 15140122 |
| Trypan Blue Stain | Invitrogen | Cat: T10282 |
| Matrigel™ Matrix | Corning | Cat: 356234 |
| Heparin | Sigma Aldrich | Cat: H3149-25KU |
| GADOVIST® (gadobutrol) | Bayer | NA |
| Fluorinert™ | 3M | Cat: 7100003808 |
| MYCi975 | Invivo Chem | Cat: V37600 CAS: 2289691-01-4 |
| InVivoMAb anti-mouse CTLA-4 (clone 9H10) | BioXcell | Cat: BE0131 |
| InVivoMAb anti-mouse PD-1 (clone RMP1-14) | BioXcell | Cat: BE0146 |
| Anti-Mouse PD-L1<br>In Vivo Antibody<br>(clone 10F.9G2) | Assay Genie | Cat: IVMB0095 |
| Anti-Mouse CD40<br>In Vivo Antibody -<br>Low Endotoxin<br>(clone FGK45) | Assay Genie | Cat: IVMB0152 |
| Anti-Mouse CD115<br>In Vivo Antibody -<br>Low Endotoxin<br>(clone AFS98) | Assay Genie | Cat: IVMB0001 |
| bAb0249 Anti-CD19<br>[6D5], Mouse IgG2a<br>mCD3e (clone 145-2C11) KIH<br>bispecific, Fc<br>Silent™, Kappa | AbsoluteAntibody | Cat: bAb0249-BS |

### Cell lines and cell culture

Murine *BCR-ABL1* B-ALL cells (PER-M60), PER-M224 (*Ts1/Cdkn2a-KRASG12D*) were developed as previously described^20,112^. Murine MLL-AF9 PER-M5 cells were obtained following established protocols^113^. Briefly, primary MLL-AF9-expressing cells were transplanted into irradiated recipient mice. MLL-AF9 PER-M5 cells were subsequently derived from secondary recipient mice that developed AML. Murine M3-9-M RMS cells were derived from *Mt1:Hgf*^Tg^;*Tp53*^+/-^ mice, which were kindly shared by Crystal Mackall, Stanford University^22^. M3-9-M cells were retrovirally transduced with MSCV-ires-pacLuc2 (kindly provided by Dr Richard Williams, St Jude Children’s Research Hospital) to drive expression of codon-optimized firefly luciferase, referred to as M3-9-M luc. Mouse MCA205 fibrosarcoma cell line was obtained from Sigma-Aldrich. Mouse WEHI164 fibrosarcoma cell line was obtained from CellBank Australia. All murine sarcoma cells were cultured in RPMI media supplemented with 10% FCS, glutamine, HEPES and penicillin/streptomycin. The murine model of G3MB expressing *Myc*^T58A^, dominant-negative *Tp53*, *GFP* and codon-optimized firefly luciferase (Myc/p53^DD^) is previously described^21^ and was generously provided by Prof. Robert Wechsler-Reya, Columbia University. The murine model of H3.1^K27M^ DMG, expressing Hist1h3b^K27M^, DNp53, Pdgfra^D842V^, Acvr1^G328V^, is previously described^24^ and was generously provided by Timothy N. Phoenix, University of Cincinnati. Myc/p53^DD^ cells were maintained and amplified by intracranial passage through C57Bl/6 mice (000664), with tumor cells harvested from donor mice on the morning of implantation. Tumors were dissociated via trituration in Neurobasal™ media, counted using Trypan Blue exclusion and resuspended in Matrigel™ prior to use.

### Mice

All animal experiments were conducted in accordance with the Australian Code for the Care and Use of Animals for Scientific Purposes and were approved by the Animal Ethics Committee of The Kids Research Institute Australia (AEC341, 367, 379, 384, 385, 386, P2106, and P2343). Mice were housed under specific and opportunistic pathogen-free conditions at The Kids Research Institute Australia Bioresources facility, maintained under a 12-hour dark/light cycle at a temperature of 20-22 °C, with access to standard chow and water *ad libitum*. Both male and female mice were used in this study, except for the M3-9-M RMS experiments, which utilized only male mice. Sample size calculations were based on data from pilot studies. Tumor implant and drug administrations were performed in the early morning and midday (ZT1 and ZT5, zeitgeber time 1; 1 hour after light onset in a 12-hour light/dark environment).

### *In vivo* tumor injection

Post-natal day (P) 1-4 pups were anesthetized by hypothermia. Hypothermic anesthesia was determined by a loss of movement in the pup and a slowing of respiratory rate. For medulloblastoma inoculation, after pups were anesthetized by hypothermia, 1 μL of cell suspension (5×10^3^ cells) in Matrigel was implanted by injecting through the skull and into the cerebellum using an ice-cold Hamilton Neuros Syringe, 0.5-1 mm lateral to the midline and 2 mm caudal of the lambdoid suture, holding the syringe at a 50-60° angle to the back of the head and injecting at a depth of 1.5-1.7 mm. Inoculation of medulloblastoma cells into the cerebellum of adult C57Bl/6 mice is previously described^114^. Briefly, animals were anesthetized using ketamine, medetomidine, and alfaxalone (70 mg/kg, 1 mg/kg, 5 mg/kg respectively, *i.p*.), or isoflurane (4% induction in air, maintained at 1.5% in air). To maintain body temperature, mice were kept on surgical warming pads or in thermal cages. Adult mice were restrained by hand, an incision was made in the scalp, and a 0.8 mm burr hole was drilled into the skull using a handheld microdrill 1 mm lateral of the midline and 2 mm caudal of the lambdoid suture. Medulloblastoma cell suspensions in Matrigel (1 μL per mouse) were implanted into the cerebellum at a depth of 2 mm using an ice-cold Hamilton Microlitre Syringe. H3.1^K27M^ DMG cells (250,000 as a 2 μL suspension in matrigel) were implanted into the brain stem of P3 pups 0.5-1 mm lateral to the midline and 2 mm caudal of the lambdoid suture, holding the syringe at a 50-60° angle to the back of the head and injecting at a depth of 2.0-2.2 mm. For subcutaneous tumors, cancer cells were suspended in PBS as 5×10^4^ or 5×10^5^ cells in 10 µL for M3-9-M RMS, MCA205, and WEHI164 or in 50% Matrigel as 5×10^5^ in 20 µL for 9464D neuroblastoma cells and pups or adult mice were inoculated in the right flank. Cells were injected using Ultra-fine 0.3 cc syringes (BD). Tumor volume (mm^3^) was measured using calipers, using the following formula: (length × width^2^)/2. For orthotopic sarcoma, M3-9-M RMS luciferase^+^ cells (1×10^3^ or 1×10^4^) were suspended in 2 µL of PBS and injected into the gastrocnemius of B6 albino or C57Bl/6 mice using 10 µL, Neuros Syringe, Model 1701 RN, 33 gauge, Point Style 4 (Cat N: HAMC65460-06 Hamilton). For leukemia, PER-M60 (1×10^4^ or 1×10^3^ cells), PER-M224 (1×10^4^ cells) or PER-M5 (1×10^5^) cells in 10 µL were injected into the facial vein of P1-2 pups or in 100 µL into the tail vein of 12-14 weeks old adult mice using ultra-fine 0.3 cc syringes (BD). Comparing leukemia progression in pups implanted at P1-2 with older mice implanted at P11 and P25 where facial and tail veins are not accessible, cells were injected via retro-orbital (RO) route. After injection, pups were recovered by warming on a surgical mat until color, movement, and respiratory rate had recovered. Upon recovery, pups were returned to the nesting dam.

### *In vivo* drug treatments

All antibodies and MYCi975 were administered by intraperitoneal injection at 5 µL/g for pups and adults. Anti-PD-L1 (10 mg/kg) or anti-PD-1 (10 mg/kg) were injected every other day for 3 doses. Anti-CTLA-4 (10 mg/kg) was given once, at the same time as anti-PD-1 for MCA205 and WEHI164 fibrosarcoma models. CD19xCD3 BiTE (1 mg/kg) was given once. MYCi975 (40 mg/kg) was dissolved in 10% DMSO, 40% PEG300, 10% Tween80 and 40% PBS, and was given daily for a maximum of 4 weeks, except in the 3 days where anti-PD-L1 was given. Anti-CD40 (5 mg/kg) was given for 2 or 3 doses, depending on the cancer model, as indicated in figures. Anti-CSF1R (40 mg/kg) was given as a single dose on day 6 or 8 post inoculation, depending on the cancer model, as indicated in figures. Pups were randomized into treatment groups so that each group contained pups from multiple different litters to account for inter-litter variation. Litter sizes of 4-8 were used in all treatment experiments.

### *Ex vivo* MRI imaging of brain tumors

Pups were transcardially perfused with 2.5 mL, and adults with 5 mL, of PBS containing 1 μg/mL Heparin and 2 mM GADOVIST® followed by 2.5 mL (pup) or 5 mL (adult) of 4% PFA/PBS containing 2 mM GADOVIST®. The whole head was collected and the bottom jaw as well as all soft tissue surgically removed from the skull. Brains (maintained within the skull) were incubated for 24 hours with gentle rocking at RT in 4% PFA/PBS containing 2 mM GADOVIST®, then stored in PBS containing 2 mM GADOVIST® and 0.02% w/v sodium azide at 4°C for 30 days. Prior to MRI acquisition, samples were soaked in Fluorinert™ FC-770 for 24 hours. MRI scans were obtained using a BioSpec 9.4T MRI (Bruker). Brains implanted with Myc/p53^DD^ cells were scanned using a 72 mm transmit-receive coil. T2-weighted 3-dimensional fast spin multi-echo scans were acquired at a resolution of 100 μm and zero-filled to a resolution of 66 μm (350 ms repetition time (TR), 30 ms echo time (TE)) at the University of Queensland, Centre for Advanced Imaging.

### *In vivo* bioluminescence imaging

Bioluminescence imaging to monitor tumor size *in vivo* was performed using a Lago X (Spectral Instruments Imaging). Mice were injected intraperitoneally with 15 mg/kg D-luciferin in DPBS prior to imaging and anesthetized with 5% isoflurane. Isoflurane anesthesia was maintained at 1.5-2% during imaging acquisition and images were acquired every minute for up to 15 minutes until the peak in photon flux was observed.

### Tissue harvest and single-cell suspensions

Tissues were collected from mice in RPMI medium containing 10% FCS and processed into single cells on the same day of collection. Unless stated otherwise in the text, subcutaneous tumors were collected 11-13 days after implant and digested using 1 mg/mL of collagenase IV, 30 U/mL of DNase I and 0.1 mg/mL of hyaluronidase. GFP^+^ Myc/p53^DD^ G3MB were collected 10-11 days after implant, and manually dissected from visible normal cerebellum using low power GFP-guided wide-field microscopy. Single cell suspensions were made as previously described^115^. Briefly, tissue was triturated 5 times in 5 mL DPBS containing calcium and magnesium (DPBS^Ca/Mg^) containing 100 U/mL collagenase IV and 10 U/mL DNase I, then incubated for 30 minutes at 37°C with gentle agitation by hand every 5 min. For scRNAseq experiments, 175 ug/mL of Liberase was added to the enzymatic digestion and 250 U/mL DNase was used. Enzymatic digestion was halted by the addition of 5 mL of ice-cold neuro-FACS buffer (2% FCS, 5 mM EDTA in Hank’s buffered salt solution (HBSS). The suspension was passed through a pre-wet 70 μm strainer, with an additional 5 mL of neuro-FACS buffer passed through to wash. Cell suspensions were centrifuged (300 x *g,* 5 min, room temperature (RT)) and erythrocytes lysed in 2 mL of red blood cell (RBC) lysis solution for 2 min at RT. Isotonicity was restored with the addition of 10 mL of DPBS and the cell suspension was centrifuged again (300 x *g*, 5 min, RT). For the leukemia PER-M60 B-ALL model, unless stated otherwise in the text, spleens and bone marrows were collected 11-12 days post-inoculation from pups, or 16-17 days from adults. These two timepoints were chosen so the leukemia burden in the bone marrow did not exceed 30% in order to have sufficient immune cells for characterization. Spleens were dissociated by passing through 70 µm strainers. For bone marrow single cell preparation, femurs and tibiae were collected, gently crushed then passed through a 70 µm strainer.

### Flow cytometry

Single-cell suspensions from various tissues were stained with fixable viability dye for 10 minutes in PBS, then washed and resuspended in staining buffer (PBS with 1% BSA and 0.01% Na azide). Cells were incubated with antibody cocktails for 30 minutes at room temperature, followed by washing. Where intracellular staining was required, cells were fixed and permeabilized using the eBioscience™ Foxp3/Transcription Factor Staining Buffer Set according to the manufacturer’s protocol. The cells were then stained with intracellular antibody mixes for 30 minutes at room temperature, washed, resuspended in staining buffer, and analyzed using a BD FACSymphony™ A5 SE Cell Analyzer. A table of antibodies is provided in Key Resource Table 1.

### T_RM_ validation using intravascular labelling

As previously described^39,40^, M3-9-M RMS or PER-M60 B-ALL bearing pups or adult mice were injected retro-orbitally with 2 µg/10 µL of PE-CD45 (clone 30F11). After 5-10 minutes, the mice were humanely euthanized. Blood and tumor tissues (M3-9-M RMS) or bone marrow (PER-M60 B-ALL) were collected and processed for flow cytometry analysis. For T_RM_ identification, cells were gated on CD8^+^ T cells that were negative for the intravascular labelling antibody (PE^-^CD45), indicating tissue residence as they were not accessible to the bloodstream at the time of antibody injection. These cells were further characterized by the absence of the lymph node homing receptor CD62L, the effector marker KLRG1, and by positive expression of CD44 and CXCR6 (M3-9-M RMS) or CD69 (PER-M60 B-ALL).

### Adoptive T cell transfer

Spleens were dissociated from adult B6 CD45.1 mice (12-14 weeks old, non-tumor-bearing). RBC lysis was performed, and CD3^+^ cells were isolated using MojoSort™ Mouse CD3 T Cell Isolation Kit. The purity of splenic T cells was verified by flow cytometry (∼95% T cells). A total of 3 million adult CD45.1 T cells were resuspended in 10 µL PBS and injected intravenously into C57BL/6J (CD45.2) pups (P0-1) through the facial vein. One-or two-days following T-cell transfer, tumor cells were introduced according to the respective model: M3-9-M RMS subcutaneously, Myc/p53^DD^ G3MB intracerebellar, or PER-M60 B-ALL intravenously. Tumors were harvested and analyzed 11–13 days after implantation. To isolate the naïve T-cell fraction, splenic cell suspensions were flow-sorted for the CD62L^⁺^CD44^low^ population; 6 × 10^5^ of these naïve T cells were then administered intravenously through the facial vein of P0-1 pups, followed by subcutaneous injection of M3-9-M RMS cells two days later. Tumors were harvested and analyzed 11 days after implantation

### Tumor mutational burden analyses

Genomic DNA was extracted from murine cancer cell lines/models using the DNeasy (Qiagen) according to the manufacturer’s protocol. Whole-genome libraries were prepared using the Illumina DNA Prep PCR-Free Library Kit and sequenced on an Illumina NovaSeq X platform at the Australian Genome Research Facility (AGRF). Variants were identifiedusing the Illumina DRAGEN pipeline (version 4.0.3) by aligning sequencing reads to strain-specific reference genomes: GRCm39 (C57BL/6J-derived) for C57BL/6 derived-samples (Myc/p53^DD^ G3MB, M3-9-M RMS, MCA205 sarcoma, PER-M60 B-ALL), and BALB_cJ_v1 for tWEHI164 sarcoma, with annotation performed using Ensembl Variant Effect Predictor (VEP)^116^. Tumor mutation burden (TMB) was calculated as the number of mutations within exonic regions divided by the total size of these regions, reported as mutations per megabase (mut/Mb).

### Single cell sample pre-processing for scRNAseq

M3-9-M RMS tumors were collected 11 days post-implant, and live single cells were isolated from tumor suspensions using fluorescence-assisted cell sorting (FACS). Myc/p53^DD^ G3MBs were collected 10-11 days post-implant and single cell suspensions were magnetically enriched for CD45^+^ cells due to the low immune infiltration in these tumors, resulting in approximately 1:1 mix of tumor and CD45^+^ cells. For PER-M60 B-ALL, bone marrow cells were collected 14 days post-implant and single cell suspensions of 1:1 mCherry^+^ (B-ALL) and mCherry^-^ cells were collected using FACS. Library construction (using the 10X Chromium 3’ workflow) and quality control was carried out by Genomics WA (M3-9-M RMS and PER-M60 B-ALL) or according to manufacturer’s protocol (Myc/p53^DD^ G3MB). Libraries were sequenced using NovaSeq S1 kits by Genomics WA.

### Mouse scRNAseq data processing

scRNA-seq data for each library were processed with CellRanger count (CellRanger V6.1.1, 10x Genomics) with a custom reference based on the mouse reference genome mm10 (Consortium 2002). Within R, gene counts per cell were merged with the phenotype data in Seurat V4^32^. Cells were filtered with a probabilistic framework using miQC^117^. For gene expression counts, individual samples were merged into one expression matrix and analyzed using the Seurat package. Transcript counts were log-normalized (Seurat, NormalizeData). SingleR^118^ was used to roughly identify cells in the combined object via known markers using several celldex mouse reference libraries. The top 2,000 most variable genes were selected using variance stabilizing transformation (FindVariableFeatures), followed by data scaling (ScaleData). Principal component analysis was then performed on this gene space (RunPCA). Clustering was carried out on the basis of the shared nearest neighbor between cells (FindNeighbors, 30 PCs) and shared nearest neighbor (SNN) clustering (FindClusters, 30 PCs, resolution of 0.5). We calculated markers for individual clusters using the FindMarkers function in Seurat (Wilcoxon’s rank sum test, Benjamini–Hochberg adjustment for multiple testing). For further analysis, we retained clusters that contained at least 50 cells. Differentially expressed genes for each phenotype on a cluster-by-cluster basis were identified using a linear model with MAST^119^. Cancer Hallmark analyses were performed on a cluster-by-cluster basis with fGSEA^120^ using the logFCs from MAST.

### RNA preparation for bulk RNA sequencing

For the Myc/p53^DD^ G3MB and M3-9-M RMS models, tumors were collected from pups and adult mice 10-12 days post-implantation, snap-frozen on dry ice, and stored at -80°C. For the MYCi947 target inhibition experiment, pediatric mice bearing M3-9-M RMS were administered MYCi975 at 40 mg/kg once daily, beginning on the second day after tumor implantation. Tumors were harvested 11 days post-implantation, prior to reaching endpoint criteria, and within 16 hours of the final dose. RNA was extracted using the RNeasy Plus Mini Kit according to the manufacturer’s instructions. For PER-M60 B-ALL model, bone marrow cells were harvested from both pups and adult mice 15 days post-implant and sorted for mCherry^+^CD19^+^ cells using FACS. RNA was extracted using the RNeasy Micro Kit following the manufacturer’s protocol. RNA was submitted to either Genomics WA (Myc/p53^DD^ G3MB and M3-9-M RMS) or AGRF (PER-M60 B-ALL) for total RNA library preparation, ribosomal RNA depletion, and sequencing.

### Mouse bulk RNAseq data processing

For initial data processing of RNAseq data all libraries were prepared using Unique Molecular Identifiers (UMIs) as paired-end reads and sequenced at Genomics WA or AGRF using the Illumina NovaSeq 6000 platform. All samples of the bulk Myc/p53^DD^ G3MB and M3-9-M RMS cohorts were processed as two read groups to achieve sufficient read depth, ranging between 35-50 million paired reads per sample. Some samples of the bulk leukemia cohort underwent additional sequencing to achieve a read depth of > 50M. Mean Phred scores for each sample, as determined by FastQC^121^ were generally high. FastQ sequences were aligned with kallisto^122^ to a custom reference based on the mouse reference genome mm10 and GENCODE M27 gene models with an index constructed of 31bp kmers (v10). The counts matrices were assembled from the kallisto output using tximport^123^, summarized at the gene and transcript levels with lengthScaledTPM. Additional annotation was obtained from BioMart^124^

Differential Expression Analysis: At the gene level, the size of the dataset was reduced by filtering out genes that had fewer than 3 genes across the samples of each phenotype. This requires a consensus of at least 3 samples amongst the animals in each group. This resulted in retention of 38777 genes for Myc/p53^DD^ G3MB, 34499 for M3-9-M RMS, and 35846 genes for PER-M60 B-ALL. Normalization was performed by variance stabilization (VST). Analysis of differential expression of genes between pup and adult samples was undertaken with DEseq2^125^. Following differential gene expression analysis, a ranked list of genes ordered by Log2FC was created and analyzed using the fGSEA package (v1.24.0)^120^ and the Molecular Signatures Database mouse hallmark gene sets using MSigDBr ^126^ (v7.5.1).

### Human bulk RNA-seq data analysis

Feature counts data generated using HTSeq for diagnostic alveolar (*n=6)* and embryonal (*n=17)* rhabdomyosarcoma tumor samples from 23 pediatric patients and 79 medulloblastoma pediatric patients at diagnosis were obtained from St. Jude Cloud^63^ and subsequently imported with httr^127^ (v1.4.6) into R^128^ (v4.2.2). Genes were filtered based on their expression level with edgeR^129^ (3.40.2) before DESeq2^125^ (v1.36.0) differential gene expression analysis. Age groups, 0-4 (rhabdomyosarcoma n=14, medulloblastoma n=37) and >14 (rhabdomyosarcoma n=9, medulloblastoma n=42), were created to obtain gene expression profiles. To control heterogeneity in these datasets, we assigned covariates RNA sequencing library preparation, disease classification and age at diagnosis into the design. Log2 gene expression RNA sequencing data for atezolizumab non-responders (PD, n=49) and responders (PR/SD, n=11) was obtained from the European Genome-Phenome Archive (EGAS00001006004)^64^, downloaded with pyEGA3 (v5.2.0) and imported into R (v4.2.2). Gene filtering and differential gene expression was performed as described above.

Gene set enrichment analysis with pre-ranked log_2_ fold change genes was performed using the MSigDBr ^126^ (v7.5.1) and clusterProfiler^130^ (v4.4.4) packages.

### Human sample alignment and data processing for scRNAseq

Human SHH-MB single-cell RNA-seq samples (n = 2) were obtained from Vladoui et al. 2019^49^. Samples were reprocessed as follows. Cell Ranger (10x Genomics, v1.2) ’count’ option with default parameters was used to filter and align sequencing reads to the hg19 reference genome build and obtain gene counts per cell. Downstream data processing was performed using Seurat (v3.2.1)^131^. Briefly, cells with a mitochondrial content higher than 2 standard deviations above the median, a number of genes of less than 400 or higher than 2 standard deviations above the median and a number of unique molecular identifiers (UMIs) higher than 2 standard deviations above the median were filtered out. Libraries were scaled to 10,000 UMIs per cell and log normalized. UMI counts and mitochondrial content were regressed from normalized gene counts and the residuals Z-scored gene-wise. Dimensionality reduction was performed using principal-component analysis applied to the top 2000 most variant genes. The first 30 principal components were then used as input for projection to two dimensions using UMAP and for clustering using a shared nearest neighbor (SNN) modularity optimization algorithm based on the Louvain algorithm on a k-nearest neighbors graph with k = 20 and a clustering resolution of 1. The cells were annotated using CORAL^132^, a pipeline for weighted consensus annotation of cell types, which uses a set of machine learning tools trained on a singlecell resolution reference. Twelve machine learning methods were used: SingleR^133^, scClassify^134^, SciBet^135^, singleCellNet^136^, scHPL^136^, linear support vector machines, Spearman correlation, scLearn^137^, ACTINN^138^, scID^139^, scAnnotate^140^, and scNym^141^. As a training reference, we used the immune cells from the scRNAseq samples from a previously reported dataset of pediatric gliomas^46^. Predicted labels from each method were aggregated into broader categories based on a pre-defined cell type ontology. Next, cells were filtered based on the expression of the PTPRC immune marker, with cells belonging to PTPRC+ clusters retained for downstream analysis. Differentially expressed genes (DEGs) were identified using the Wilcoxon rank sum test as implemented in the FindMarkers function from the Seurat package, subsampling cells as needed to obtain balanced groups for comparison (logfc.threshold = 0, min.pct = 0, max.cell.per.ident = 61 for macrophages and 75 for myeloid). Significant genes were defined based on fold change and percentage of cells expressing the genes (log2FC > 1, p_val_adj < 0.05, pct.1 and pct.2 > 0.001). Gene set enrichment analysis for DEGs ranked by average log2 fold change was performed using the GSEA function of the clusterProfiler package (v. 4.6.2) and the Hallmarks gene sets (MSigDBr v. 7.5.1). For the Group 3/4 medulloblastoma patient single cell analysis, data were downloaded from the GEO (GSE155446) and analyzed using Seurat V4. Patients with a diagnosis of Group 3 or Group 4 medulloblastoma as per the original publication^65^ were grouped into two age groups for analysis, ≤ 4 years and ≥ 10 years of age (n = 6 per group). Differentially expressed genes for each age group on a cluster-by-cluster basis were identified using a linear model with MAST^119^ on clusters identified in the original publication. Gene set enrichment analysis was performed on a cluster-by-cluster basis using the fGSEA package (v1.24.0)^120^ and the Molecular Signatures Database mouse hallmark gene sets using MSigDBr (v7.5.1), based on the log2FCs calculated by MAST.

Pre-processed and annotated scRNAseq data (processed as previously described^111^) were acquired from the Alex’s Lemonade Stand Single-cell Pediatric Cancer Atlas (ALS ScPCA) for the projects SCPCP000007 (AML) and SCPCP000008 (B-ALL)^47,48^. Projects were included on the basis of the following selection criteria: a) the inclusion of at least 3 patients under 4 years of age and 3 patients over 14 years of age, b) who received the same cancer type diagnoses, and c) with samples collected from the same tissue/organ (to account for microenvironmental impacts on immune populations). All available samples that met the criteria were analyzed from SCPCP0000007 (AML, n=4 and n=8 of <4 and >14 year old patients, respectively) and n=12 of the youngest and oldest patients per age group (<4 and >14) from SCPCP000008 (B-ALL) were analyzed. SingleCellExperiment objects were converted into Seurat objects and downstream analysis was carried out in Seurat V4. Per individual SCPCP project, Seurat objects were integrated using the top 2,000 most variable genes using Seurat integration functions (FindVariableFeatures, SelectIntegrationFeatures, FindIntegrationAnchors, IntegrateData). Data was scaled (ScaleData), and principal component analysis was performed (RunPCA). Using the top 30 principal components (PCs), UMAP dimensionality reduction was carried out, followed by clustering on the basis of shared nearest neighbors (FindNeighbors, 30 PCs and FindClusters, resolution = 0.5). The existing reference-based and marker-gene-based annotations from the Alex’s Lemonade Stand ScPCA pipeline were used to identify malignant and immune populations. Differentially expressed genes for each phenotype on a cluster-by-cluster basis were identified using a linear model with MAST^119^. Cancer Hallmark analyses were performed on a cluster-by-cluster basis with fGSEA^120^ using the logFCs from MAST.

### Statistical analysis

To investigate whether age impacted tumor growth rate, we inoculated mice from P3-P119 with M3-9-M RMS tumors (n=46 total) and used a linear mixed-effects model with random intercepts for individual mice. Using tumor volume measurements (log transformed) as the dependent variable, time (post-inoculation (days) and age (at tumor inoculation) were included as fixed effects. An interaction term between time and age was added to evaluate whether tumor growth rate over time changed with increasing age. The lme4 package (v. 1.1-34)^142^ was used, with the model specified as log(tumor size) ∼ time * age + (1|MouseID). For all other statistical analyses, GraphPad Prism 9 software was used. Significance was calculated using unpaired Student’s t-test when comparing difference between two groups, mixed-effects models for comparing growth curves between two groups and Log-rank (Mantel-Cox) for survival analysis, or as indicated in figure legends. Data are shown as the mean ± SEM, and statistical significance is shown as * p < 0.05, ** p < 0.01, *** p < 0.001, and **** p < 0.0001.

