## Supplemental data for "Age-dependent tumor-immune interactions underlie immunotherapy response in pediatric cancer"

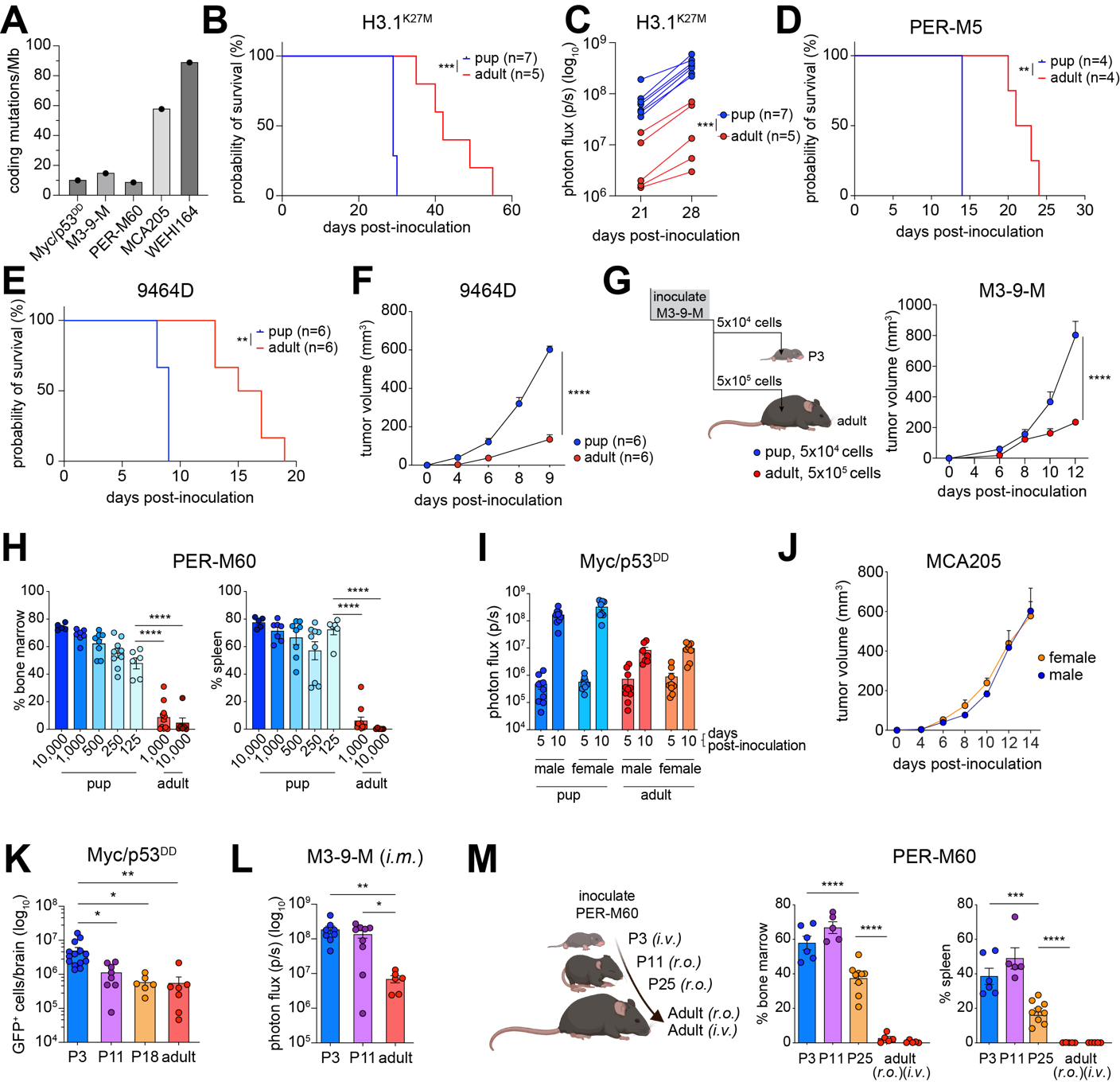

**Figure S1. Pediatric microenvironment promotes tumor growth. Related to Figure 1.** (A) Coding mutation frequencies across several cancer cell lines used for murine model establishment. (B) Survival of pup and adult mice bearing H3.1^K27M^ DMG. (C) Quantified bioluminescent flux of H3.1^K27M^ DMG tumors in pup or adult mice 21- and 28-days post-inoculation. (D) Survival of pup (n = 4) and adult mice (n = 4) bearing PER-M5 AML. (E) Survival of pup (n = 6) and adult mice (n = 6) bearing 9464D neuroblastoma cells. (F) Tumor volumes of 9464D neuroblastoma cells implanted subcutaneously in the right flank of pup (n=6) and adult mice (n=6). (G) Experiment schematic where P3 mice are inoculated with ten-fold fewer M3-9-M RMS cells than adult mice (*left*) and tumor growth (*right*). (H) Percentage of PER-M60 B-ALL cells in bone marrow (*left*) and spleen (*right*) following intravenous inoculation of 125–10,000 cells in pediatric mice and 1,000–10,000 cells in adult mice. (I) Bioluminescent photon flux of Myc/p53^DD^ G3MB tumor-bearing pups and adults at the indicated days post-implant, split by sex (n=9-10/group). (J) Tumor growth of MCA205 fibrosarcoma in male versus female pups (n=5-6/group). (K) Enumeration of GFP^+^ Myc/p53^DD^ G3MB cells in the whole brain of pups implanted at P3, P11, P18 and adult mice (n≥6). (L) Quantified bioluminescent flux of M3-9-M RMS cells in the gastrocnemius 10 days post-implantation in pups implanted at P3, P11 and adult mice (n≥6). (M) Experiment schematic (left) and percentage of PER-M60 B-ALL cells in bone marrow (*middle*) and spleen (*right*) following intravenous inoculation of 1000 cells in pups at P3, P11, P25 and adult mice (n≥6). *i.m*: intramuscular, *i.v*: intravenous, *r.o*. retro-orbital. Data are presented as mean ± SEM. Significance was calculated using Log-rank (Mantel-Cox) test (B,D,E), two-way ANOVA (C), mixed-effects models (F,G), or one-way ANOVA with Šídák’s multiple comparisons (K-M). * p < 0.05, ** p <0.01, *** p < 0.001, **** p < 0.0001.

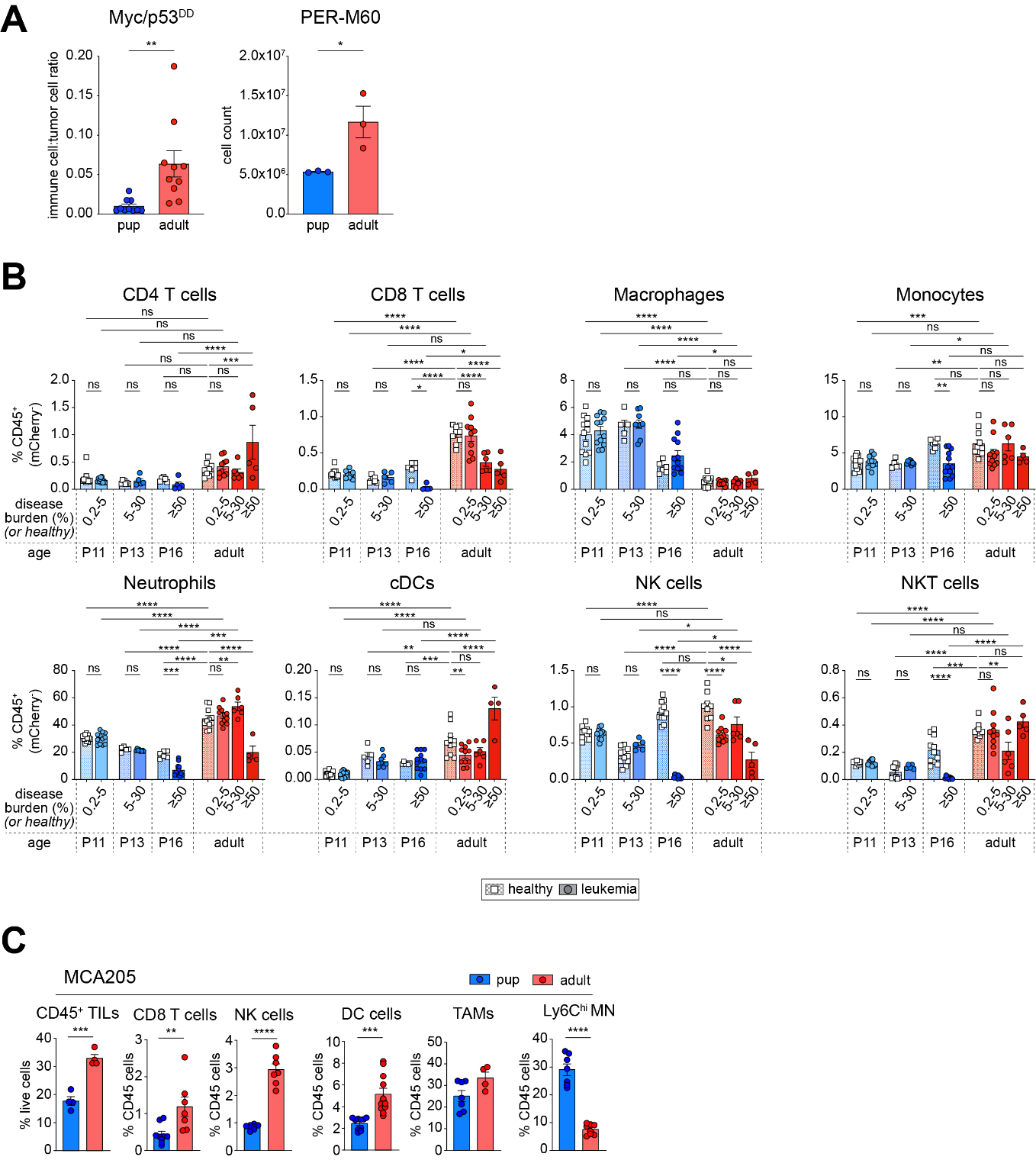

**Figure S2. Age-dependent differences in immune cell frequency across murine cancer models and under healthy conditions, related to figure 2.** (A) Quantification of age-dependent differences in TILs from pediatric and adult mice bearing Myc/p53^DD^ G3MB tumors or PER-M60 B-ALL. Data shown as the immune infiltration ratio (ratio of CD45⁺ immune cells to GFP⁺ tumor cells, *left*) in Myc/p53^DD^ G3MB tumors, and as absolute numbers of CD45⁺mCherry⁻ immune cells in the tibia bone marrow from mice bearing PER-M60 B-ALL (*right*), unpaired Student’s t-test p values indicated. (B) Bar graphs depicting the abundance of the indicated immune populations as a proportion of the total (non-malignant, mCherry^-^) CD45^+^ immune population in healthy controls *(white squares)* and age-matched leukemia-bearing *(filled circles)* pediatric *(blues)* and adult *(reds)* mice. Šídák's adjusted p values are indicated. (C) Frequency of total TILs (as a percentage of total live cells, *far left*) and composition of distinct immune subsets (expressed as a percentage of total TILs) in MCA205 fibrosarcoma tumors grown in pediatric and adult mice, unpaired Student’s t-test p values indicated. Data is presented as mean ± SEM. * p <0.05, ** p<0.01, *** p<0.001, **** p<0.0001.

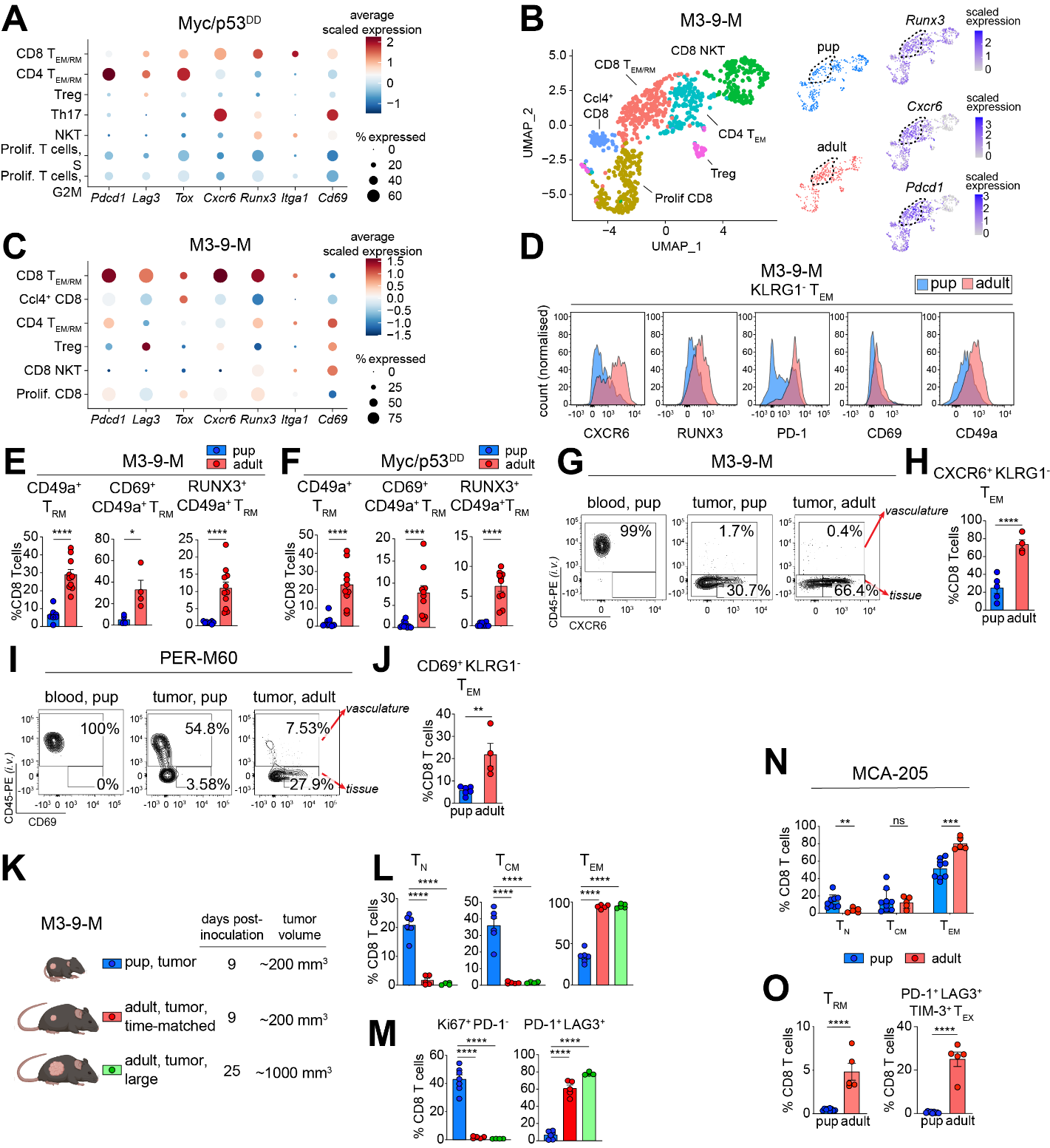

**Figure S3. Phenotypic differences in CD8⁺ T cells within tumors from pediatric and adult mice, related to figure 2**. (A) Balloon plots of the average scaled expression of the indicated T_RM_ and T_EX_ related genes across T cell scRNAseq subpopulations in Myc/p53^DD^ G3MB tumors shown in Figure 2D_._ (B) Unsupervised UMAP of scRNAseq data showing T cells from M3-9-M RMS grown in pups and adult mice *(left, middle)*, and the expression pattern of *Runx3*, *Cxcr6, and Pdcd1 (right)*. (C) Balloon plots of the average scaled expression of the indicated T_RM_ and T_EX_ related genes across T cell scRNAseq subpopulations in M3-9-M RMS tumors shown in (B). (D) Representative flow cytometry histograms of different T_RM_ associated markers in M3-9-M RMS tumors and their frequency within (E) M3-9-M RMS or (F) Myc/p53^DD^ G3MB tumors growing in pup and adult mice. (G) Representative flow cytometry plots of CD8^+^ T cells in the blood and tumor tissue after intravascular labelling using CD45-PE antibody and (H) frequency of PE^-^/CXCR6^+^/KLRG1^-^ CD8^+^ T cells in M3-9-M RMS tumors growing in pup and adult mice. (I) Representative flow cytometry plots of CD8^+^ T cells in the blood and bone marrow after intravascular labelling using CD45-PE antibody and (J) frequency of PE^-^/CD69^+^/KLRG1^-^ CD8^+^ T cells in PER-M60 B-ALL growing in pups and adult mice. (K) Experimental schema for L-M. Quantitation of (L) T_N_, T_CM_, T_EM_, and (M) Ki67^+^PD-1^-^ and PD-1^+^LAG3^+^ cells as a proportion of total CD8^+^ T cells in M3-9-M RMS tumors from pup or adult mice with small or large tumors. (N-O) Quantitation of (N) T_N_, T_CM_, T_EM_, and (O) T_RM_ and PD-1^+^LAG3^+^TIM3^+^ T_EX_ cells as a proportion of total CD8^+^ T cells in MCA205 tumors from pup or adult mice. Data is presented as mean ± SEM. * p <0.05, ** p<0.01, *** p<0.001, **** p<0.0001.

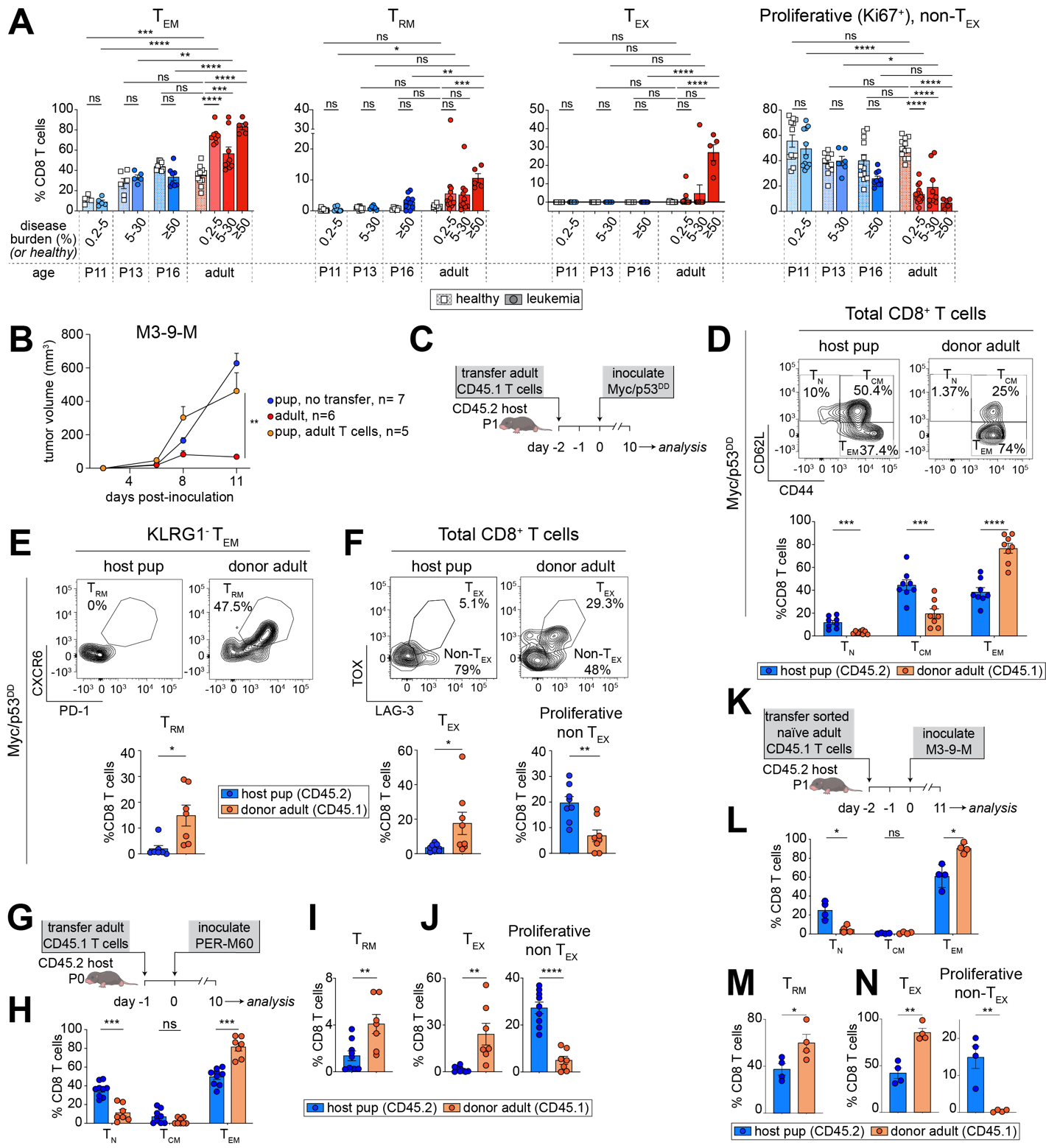

**Figure S4. Developmental origin shapes the phenotype of Intratumoral CD8⁺ T cells, related to figure 2**. (A) Frequency of T_EM_, T_RM_, T_EX_, proliferative (Ki67^+^) non-T_EX_ in CD8^+^ T cells in healthy controls *(white squares)* and age-matched leukemia-bearing *(filled circles)* pediatric *(blues)* and adult *(reds)* mice. Šídák's adjusted p values are indicated. (B) Tumor volume of M3-9-M RMS in pups after they received 3×10^6^ adult CD3^+^/CD45.1^+^ cells 2 days before M3-9-M inoculation, compared to adult mice. (C) Experimental schema for D-F. (D-F) Representative flow cytometry plots *(top)* and frequency *(bottom)* of (D) T_N_, T_CM_, T_EM_; (E) T_RM_; and (F) T_EX_ and proliferative non-T_EX_ in pup host and donor adult CD8^+^ T cells in Myc/p53^DD^ tumors. (G) Experimental schema for H-J. (H-J) Frequency of (H) T_N_, T_CM_, T_EM_; (I) T_RM_; and (J) T_EX_ and proliferative non-T_EX_ in pup host and donor adult CD8^+^ T cells in PER-M60 B-ALL bone marrow. (K) Experimental schema for adoptive transfer of purified naïve adult CD8⁺ T cells (CD62L⁺CD44^low^) into pediatric mice prior to implanting with M3-9-M RMS cells. (L-N) Frequency of (L) T_N_, T_CM_, T_EM_; (M) T_RM_; and (N) T_EX_ and proliferative non-T_EX_ in pup host and donor adult CD8^+^ T cells in M3-9-M RMS. Data presented as mean ± SEM, n ≥ 4 mice per group. Statistical significance was determined by one-way ANOVA with Šídák’s multiple comparisons test (A), mixed-effects model (B), unpaired Student’s t-test with Holm-Šídák correction (D, H, L), or unpaired Student’s t-test (E, F, I, J, M, N). * p < 0.05, ** p < 0.01, *** p < 0.001, and **** p < 0.0001.

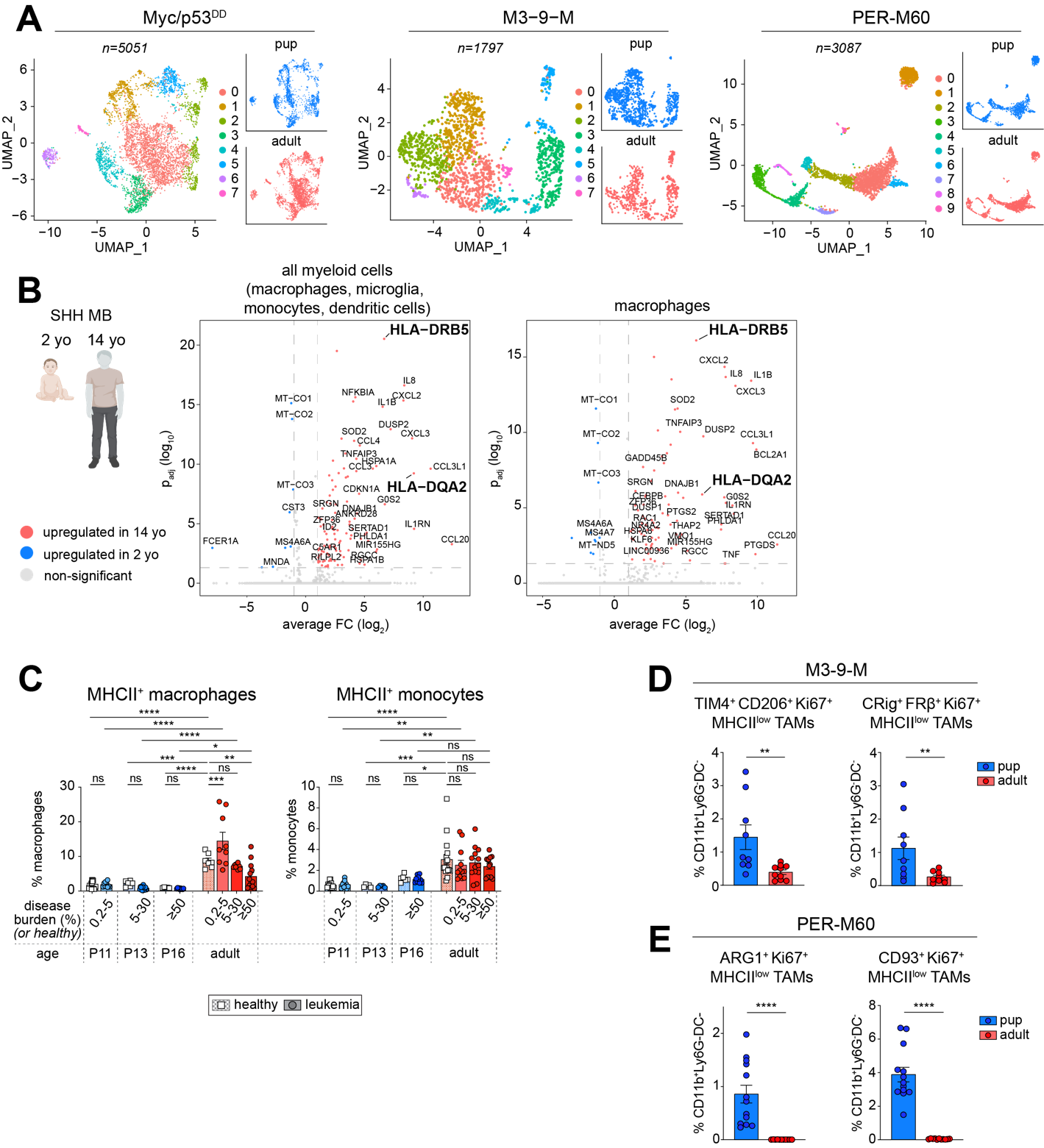

**Figure S5. TAMs and monocytes in pediatric mice tumours and lower MHCII expression in myeloid cells in an infant with SHH MB., related to figure 3**. (A) UMAP of scRNAseq data from Myc/p53^DD^ G3MB tumors, M3-9-M RMS and PER-M60 B-ALL bone marrow showing sub-clustering of TAMs and monocytes populations. Clusters identification is provided in Supplemental Table 1. (B) Volcano plot of differentially expressed genes in all myeloid cells and macrophages from two SHH MB patients of different ages. (C) Frequency of MHCII^+^ macrophages *(left)* and MHCII^+^ monocytes *(right)* in healthy controls *(white squares)* and age-matched leukemia-bearing *(filled circles)* pediatric *(blues)* and adult *(reds)* mice. Šídák's adjusted p values are indicated. (D) Frequencies of TIM4^+^CD206^+^Ki67^+^MHCII^low^ TAMs (*left*) and CRig^+^FRβ^+^Ki67^+^MHCII^low^ TAMs (*right*) as percentage of parent population (CD11b^+^/Ly6G^-^ DC^-^) in M3-9-M RMS tumors in pediatric and adult mice. (E) Frequencies of ARG1^+^/Ki67^+^/MHCII^low^ TAMs (*left*) and CD93^+^/Ki67^+^/MHCII^low^ TAMs (*right*) as percentage of parent population (CD11b^+^/Ly6G^-^DC^-^) in the bone marrow of PER-M60 B-ALL bearing pediatric and adult mice. Sample sizes: n = 3-4 per group (A), n = 1 per group (B), n ≥ 5 per group (C) and n ≥ 8 per group (D,E). Significance was calculated using unpaired Student’s t-tests (D,E). * p < 0.05, ** p < 0.01, *** p < 0.001, **** p < 0.0001

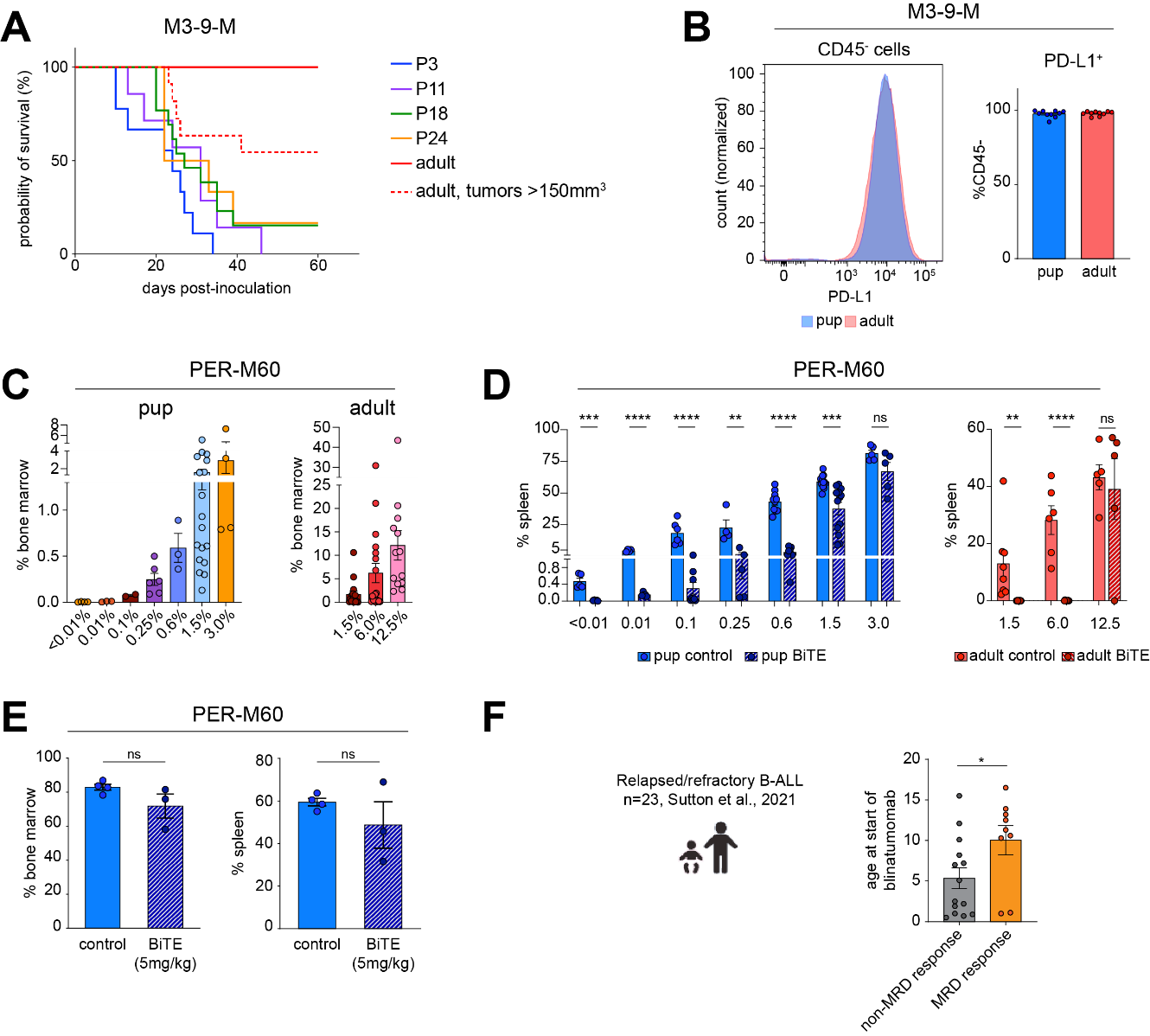

**Figure S6. Responses to immunotherapy in mice with RMS or B-ALL. Related to Figure 4.** (A) Survival curves of pediatric and adult mice inoculated with M3-9-M RMS cells and treated with anti-PD-L1 antibody, demonstrating significantly improved efficacy in adults (corresponding untreated controls shown in Figure 1K). (B) Representative flow cytometry histogram indicating PD-L1 expression (*left*) on CD45^-^ cells within M3-9-M RMS tumors obtained from pediatric (*blue*) and adult (*red*) mice 12 days after implant and frequency of positive cells (*right*, mean ± SEM, n≥10 from at least 2 independent experiments). (C) Percentage of PER-M60 B-ALL cells in bone marrow from pediatric (*left*) and adult (*right*) mice. Data represent baseline leukemia burden measurements (mean percentage, n ≥ 2 mice per group) performed prior to randomization for subsequent BiTE treatment in Figure 4H. (D) Percentage of PER-M60 B-ALL in the spleen after CD19×CD3 BiTE treatment in pediatric (*left*) and adult mice (*right*), matched data to bone marrow data presented in Figure 4H. (E) Percentage of PER-M60 B-ALL cells of total cells from bone marrow (*left*) or spleen (*right*) in pediatric mice 5 days following treatment with vehicle (n = 4) or 5 mg/kg CD19×CD3 BiTE (n = 3). (F) Age and response to blinatumomab in pediatric B-ALL patients. Pediatric B-ALL patients minimal residual disease response (MRD, x-axis) versus age of patients at start of blinatumomab treatment (y-axis), n=23. Data presented as mean ± SEM. Sample sizes: n ≥ 7 per group (A), n=10 per group (B), or n ≥ 3 per group (D-E). Statistical significance was determined using an unpaired Student’s t-tests (D-F). * p < 0.05, ** p < 0.01, *** p < 0.001, **** p < 0.0001.

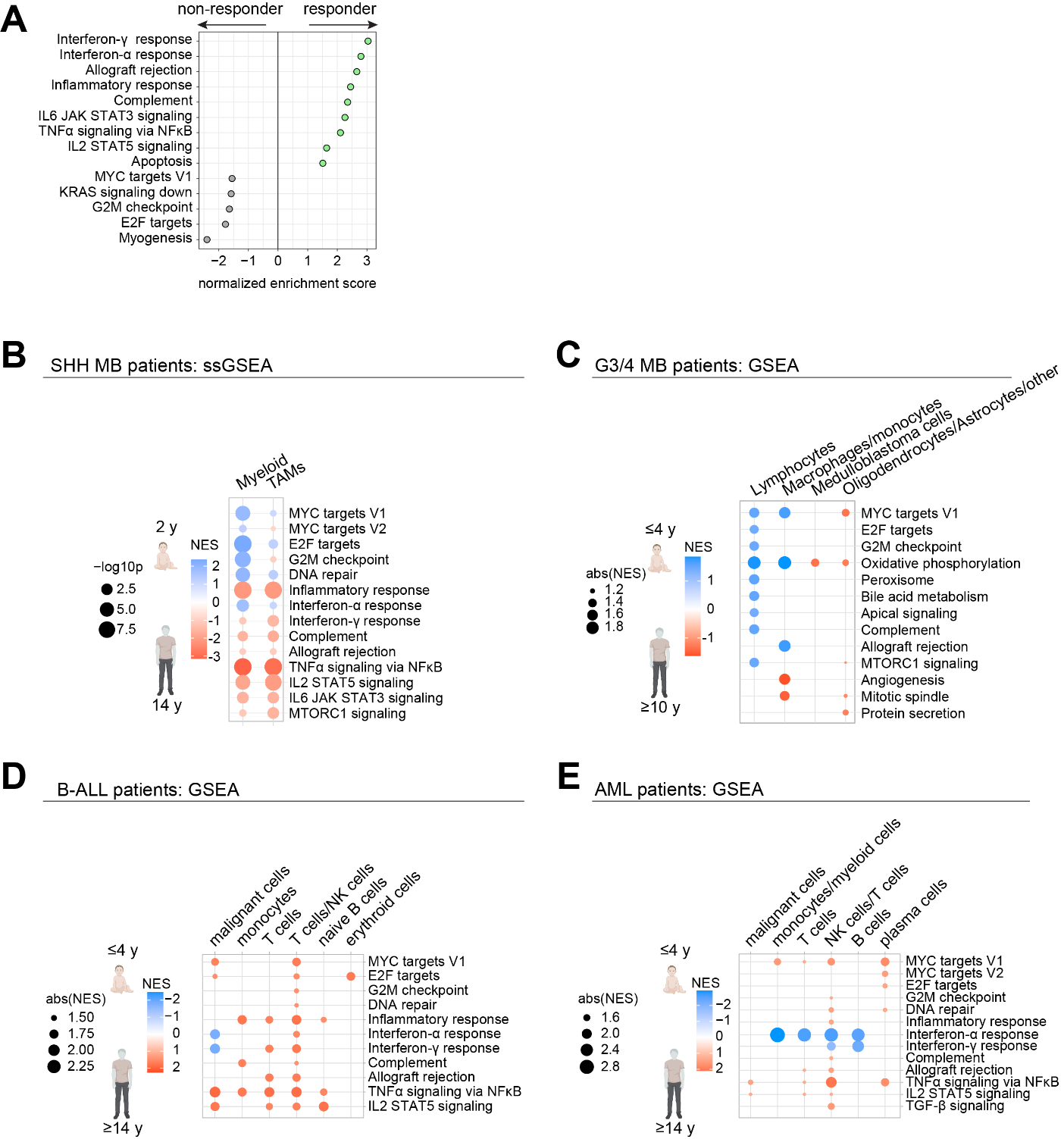

**Figure S7. Age‑dependent enrichment of MYC target and immune gene signatures in pediatric cancer patients. Related to Figure 5.** (A) GSEA of bulk tumor RNAseq from pediatric cancer patients stratified by clinical response to anti-PD‑L1 therapy (responders vs non‑responders; n=60). (B) ssGSEA of myeloid cells from an infant (2 years old) and an adolescent (14 years old) with SHH MB. (C-E) GSEA comparing gene expression in the indicated immune cells from patients with (B) Group 3/4 medulloblastoma (G3/4 MB), aged 0–4 years compared to those over 10 years old (n = 6/group). (C) B-ALL patients, aged 0-4 compared to those over 14 years old (n = 6/group) and (D) AML patients, aged 0-4 (n=4) compared to those over 14 years old (n=8). All enrichments shown reach statistical significance (adjusted p < 0.05).

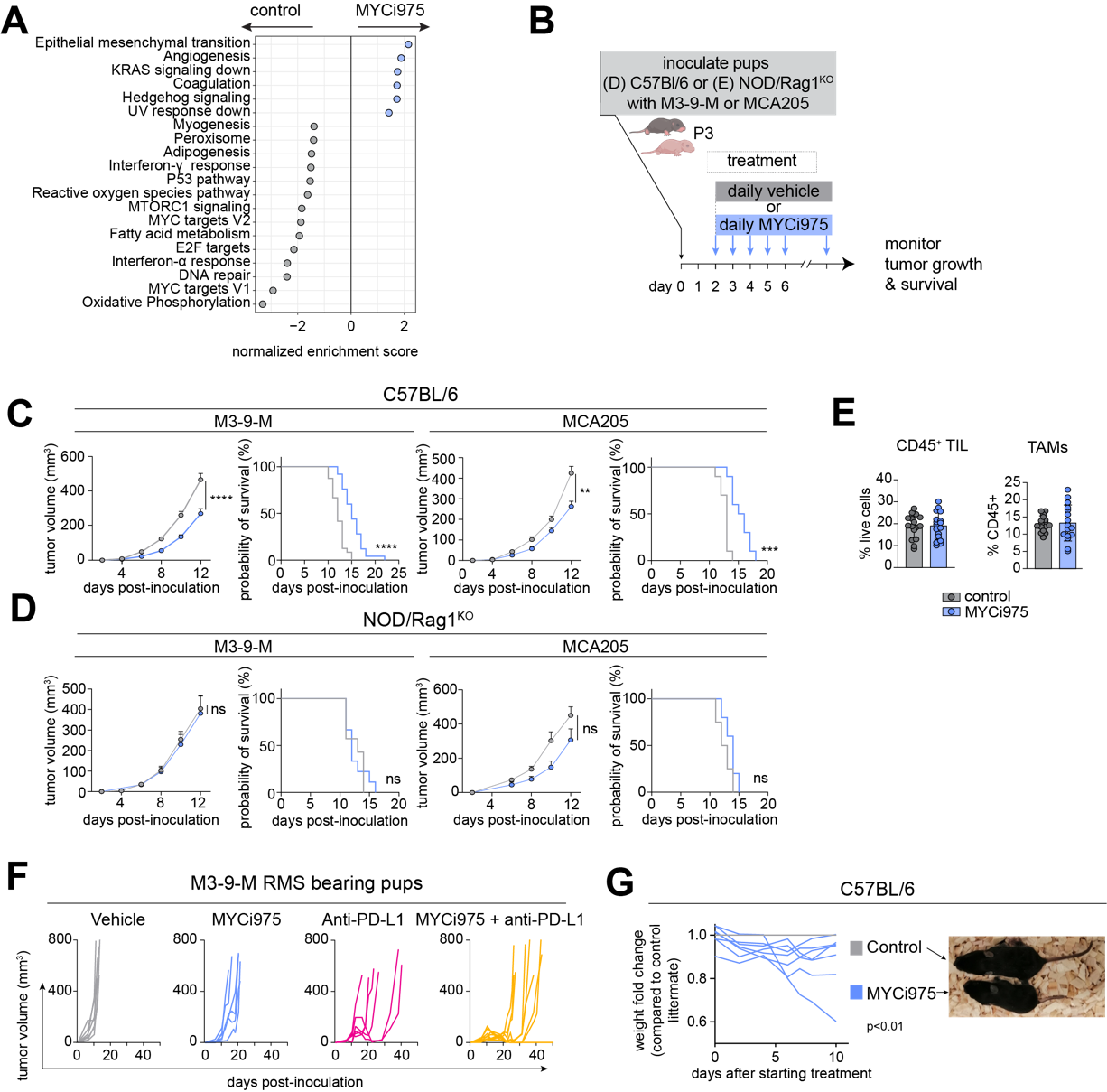

**Figure S8. MYC inhibition enhances ICT response in pediatric mice. Related to Figure 5.** (A) GSEA of bulk M3-9-M RMS tumor RNAseq from MYCi975-treated or control pediatric mice, n=8 per group. (B) Schematic for C-D. (C-D) Tumor growth and survival curves of pups bearing M3-9-M RMS (*left* *panels*) or MCA205 fibrosarcoma (*right* *panels*) in (C) immunocompetent C57Bl/6 or (D) immunocompromised NOD/Rag1^KO^ pediatric mice following treatment with vehicle control (*grey*) or MYCi975 (*pale blue*). (E) Frequency of TILs as percentage of live cells and abundance of TAMs as percentage of TILs in pediatric mice-bearing M3-9-M RMS following treatment with vehicle control (*grey*) or MYCi975 (*pale blue*). (F) Individual tumor growth curves of pups bearing M3-9-M RMS after treatment with vehicle control (*grey*), MYCi975 (*pale blue*), anti-PD-L1 (*pink*), or a combination of MYCi975 and anti-PD-L1 (*yellow*). (G) Fold change in body weight of pups treated with MYCi975 (*pale blue*) compared to vehicle-treated littermates (*grey*), and representative picture of a vehicle and a MYCi975-treated pup 8 days after starting treatment. Data are represented as mean ± SEM. Sample sizes: n ≥ 10 per group (C-D; except MCA205 in NOD/Rag1 mice, n = 4 per group), n=18 per group (E), and n ≥ 6 per group (F-G). Statistical significance was calculated using mixed-effects models (growth curves in C,D,G) and the log-rank (Mantel-Cox) test (survival curves in C-D). ns = not significant, * p < 0.05, ** p < 0.01, *** p < 0.001, or **** p < 0.0001.

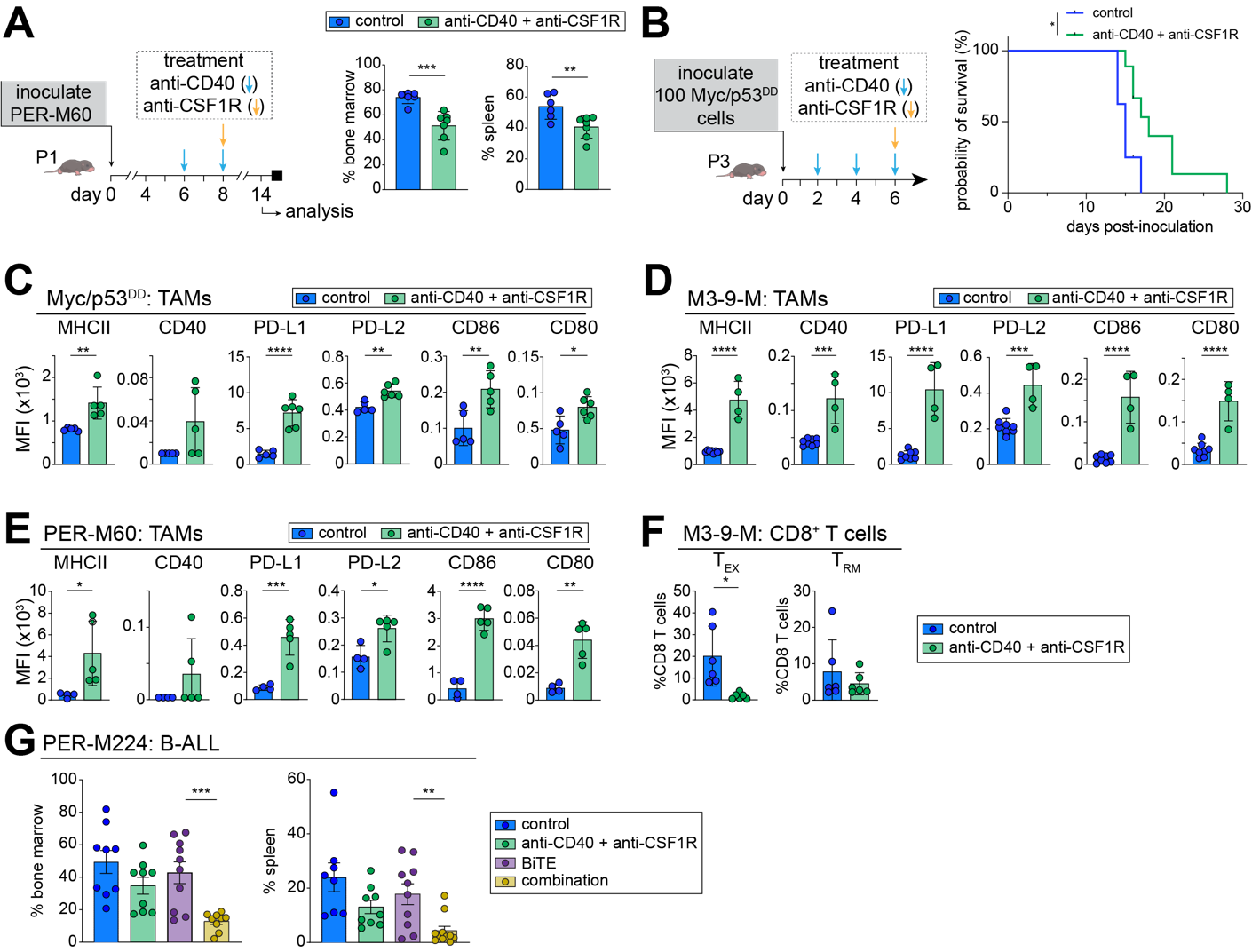

**Figure S9. Anti-CD40/CSF1R reshapes the pediatric TIME. Related to Figure 6.** (A) Treatment schema (*left*) and percentage of PER-M60 B-ALL cells in the bone marrow and spleen of pediatric mice treated with anti-CD40/CSF1R (*right*). (B) Treatment schema (*left*) and survival of mice with Myc/p53^DD^ G3MB treated with anti-CD40/CSF1R (*right*). (C-E) Median fluorescence intensity (MFI) of MHCII, CD40, PD-L1, PD-L2, CD86 and CD80 in TAMs in (C) Myc/p53^DD^ G3MB, (D) M3-9-M RMS, and (E) PER-M60 B-ALL related to the data presented in Figure 6C-G. (F) Frequency of T_EX_ and T_RM_ in intratumoral CD8^+^ T cells from M3-9-M RMS bearing pediatric mice treated with anti-CD40/CSF1R, compared to isotype control treated mice. (G) Percentage of B-ALL in the bone marrow and spleen of pediatric mice bearing PER-M224 B-ALL and treated with vehicle, BiTE, anti-CD40/CSF1R or the combination. Data are represented as mean ± SEM (A, C-G). Sample sizes: n ≥ 4 per group (A–F), n ≥ 8 per group (G). Statistical significance was calculated using mixed-effects models and unpaired Student's t-test (A and C–G), and the log-rank (Mantel-Cox) test for survival curves (B). Statistical significance is indicated as * p < 0.05, ** p < 0.01, *** p < 0.001, or *** p < 0.0001.

**Additional information**

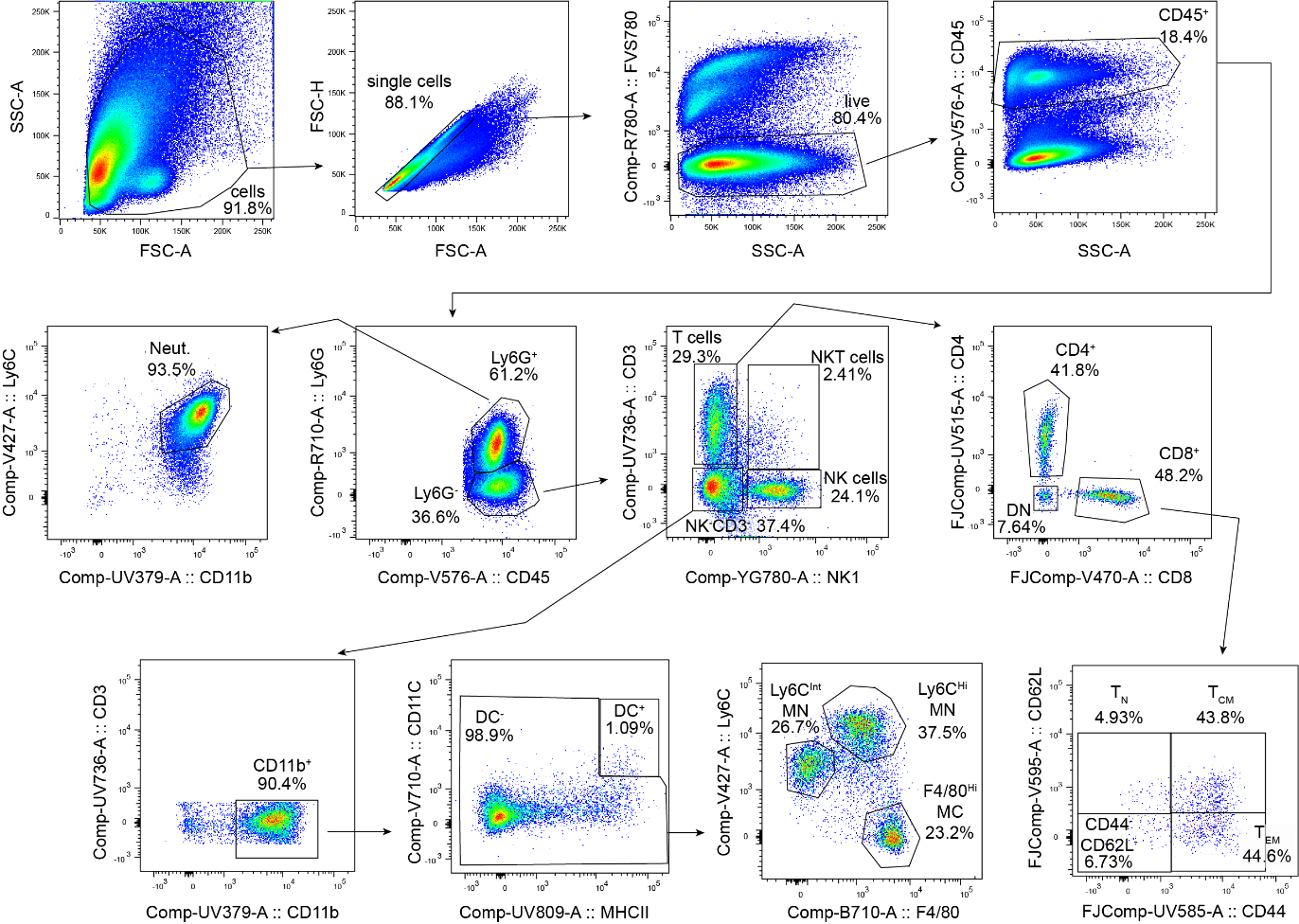

**Gating strategy**
